# Epigenetic variation impacts ancestry-associated differences in the transcriptional response to influenza infection

**DOI:** 10.1101/2022.05.10.491413

**Authors:** Katherine A Aracena, Yen-Lung Lin, Kaixuan Luo, Alain Pacis, Saideep Gona, Zepeng Mu, Vania Yotova, Renata Sindeaux, Albena Pramatarova, Marie-Michelle Simon, Xun Chen, Cristian Groza, David Lougheed, Romain Gregoire, David Brownlee, Yang Li, Xin He, David Bujold, Tomi Pastinen, Guillaume Bourque, Luis B Barreiro

## Abstract

Humans display remarkable inter-individual variation in immune response when exposed to identical immune challenges. Yet, our understanding of the genetic and epigenetic factors contributing to such variation remains limited. Here we carried out in-depth genetic, epigenetic, and transcriptional profiling on primary macrophages derived from a panel of European and African-ancestry individuals before and after infection with influenza A virus (IAV). We show that baseline epigenetic profiles are strongly predictive of the transcriptional response to IAV across individuals, and that ancestry-associated differences in gene expression are tightly coupled with variation in enhancer activity. Quantitative trait locus (QTL) mapping revealed highly coordinated genetic effects on gene regulation with many cis-acting genetic variants impacting concomitantly gene expression and multiple epigenetic marks. These data reveal that ancestry-associated differences in the epigenetic landscape are genetically controlled, even more so than variation in gene expression. Lastly, we show that among QTL variants that colocalized with immune-disease loci, only 7% were gene expression QTL, the remaining corresponding to genetic variants that impact one or more epigenetic marks, which stresses the importance of considering molecular phenotypes beyond gene expression in disease-focused studies.

## Introduction

Inter-individual differences in the transcriptional response of innate immune cells to infectious agents are common and likely contribute to varying susceptibility to infectious diseases, inflammation, and autoimmune disorders (Brinkworth and Barreiro 2014; Duffy et al. 2014; Pennington et al. 2009). Although a substantial fraction of transcriptional heterogeneity in the response to infection is likely attributable to environmental factors, several studies have shown that host genetics also plays an important role (Bakker et al. 2018; Nédélec et al. 2016; Piasecka et al. 2018; Quach et al. 2016; Randolph et al. 2021). For example, it has been shown that ∼30% of the transcriptional differences between European and African ancestry individuals in their immune responses to influenza A infection can be explained by expression quantitative trait loci (eQTL) that vary in allele frequency across populations (Randolph et al. 2021). Similar genetic contributions to ancestry-associated differences in the transcriptional response to intracellular bacterial pathogens and immune stimuli have been reported. (Nédélec et al. 2016; Barreiro et al. 2012; Quach et al. 2016).

However, much of the variance in immune responses observed at the population level remains unexplained by genetics alone (Bakker et al. 2018; Piasecka et al. 2018; Aguirre-Gamboa et al. 2016; Li et al. 2016). Other factors that have been linked to variation in immune responses include sex, age (Bakker et al. 2018; Piasecka et al. 2018), gut microbiome diversity (Schirmer et al. 2016), and the social environment (Snyder-Mackler et al. 2016, 2020; Cole 2014). Although less studied, epigenetic variation is also likely to play an important role in explaining immune response variance. The most well studied epigenetic responses to immune stimuli involve the post-translational modification of histone tails at promoter and enhancer regions (Bierne et al. 2012; Monticelli and Natoli 2013). Histone acetylation is strongly associated with the activation of many pro-inflammatory genes (Ghisletti et al. 2010; Qiao et al. 2013), whereas histone deacetylation is often associated with gene repression in the context of inflammation (Villagra et al. 2009). Moreover, certain inflammatory signals (e.g., β-glucan or Bacillus Calmette–Guerin (BCG) vaccination) or even lifestyle factors (e.g., diet) are thought to be able to “educate” the chromatin state of innate immune cells, notably monocytes/macrophages, resulting in a stronger transcriptional response during reinfection (Bekkering et al. 2021; Zhang and Cao 2021). This suggests environmentally-induced epigenetic changes may represent crucial determinants of an individual’s ability to respond to pathogens.

Although the term epigenetics means “above the genetics”, genetic variation has also been shown to play a substantial role in the degree of epigenetic variation across individuals (Chen et al. 2016; Degner et al. 2012; Carja et al. 2017; Husquin et al. 2018.; Kasowski et al. 2013; McVicker et al. 2013; Waszak et al. 2015). In human lymphoblastoid cell lines, genetic variation has been shown to impact the levels of chromatin accessibility at thousands of enhancer and promoter elements throughout the genome (Degner et al. 2012). Likewise, genetically controlled variation in chromatin accessibility has been shown to impact the magnitude of the response engaged by human macrophages in response to *Salmonella* (Alasoo et al. 2018). Thus, it is likely that variation in epigenetic profiles across individuals and populations – whether genetically controlled or not – can ultimately represent a key contributor to population variation in innate immune responses and susceptibility to disease. However, despite intense efforts to generate comprehensive epigenomic atlases across many tissues and cell types (The ENCODE Project Consortium et al. 2007, 2012; Roadmap Epigenomics Consortium et al. 2015; Fernández et al. 2016), there are no comprehensive maps of population level variation in epigenetic levels in primary innate immune cells before and after infection, preventing the formal evaluation of such hypotheses.

To address this gap, we carried out an in-depth genetic and epigenetic characterization of primary macrophages derived from 35 individuals with varying degrees of European and African ancestry at both baseline and after infection with influenza A. The data generated herein helps fill a critical gap in biomedical research: the lack of non-European ancestry individuals among cohorts designed to study immune variance in the general population and in genomic studies more generally. All data generated in this study are freely accessible via a custom web-based browser that enables easy querying and visualization of all the data generated (https://computationalgenomics.ca/tools/epivar).

## Results

### Transcriptional and epigenetic response to influenza infection

We infected monocyte-derived macrophages (MDMs) derived from a diverse panel of 35 healthy individuals with influenza A virus (IAV), commonly known as flu. We focused on macrophages as they are the primary source of type I interferon (IFN) and pro-inflammatory cytokines during flu infection, and therefore play a central role in viral clearance and the regulation of the pathology during infection (Meischel et al. 2020; Ichinohe, Pang, and Iwasaki 2010; Diebold et al. 2004). Following 24-hours of flu infection, we collected from matched non-infected (NI) and infected samples data on *(i)* gene expression (RNA sequencing), *(ii)* chromatin accessibility (assay for Transposase-Accessible Chromatin using sequencing; ATAC-seq), *(iii)* levels of histone marks associated with promoters (H3K4 trimethylation, or H3K4me3), enhancers (H3K4 monomethylation, or H3K4me1), and their activation levels (H3K27 acetylation, or H3K27ac), as well as a general repressive mark (H3K27 trimethylation, or H3K27me3), and *(iv)* methylation levels (as measured by whole genome bisulfite sequencing; WGBS) (Fig. 1A). In addition, we identified genetic variants for each individual using high-coverage (30X) whole genome sequencing. In total, we obtained over 211 billion reads across the different assays, generating the most extensive dataset to date of the combined transcriptional and epigenetic response to flu at the population level (Table S1). All assay-specific quality control metrics, including percentage of mapped reads, number of CpG sites covered per sample, or the fraction of all mapped reads that fall into the called peak regions (i.e., FRIP scores) indicate that the data is of high quality (Table S1).

**Figure 1.**
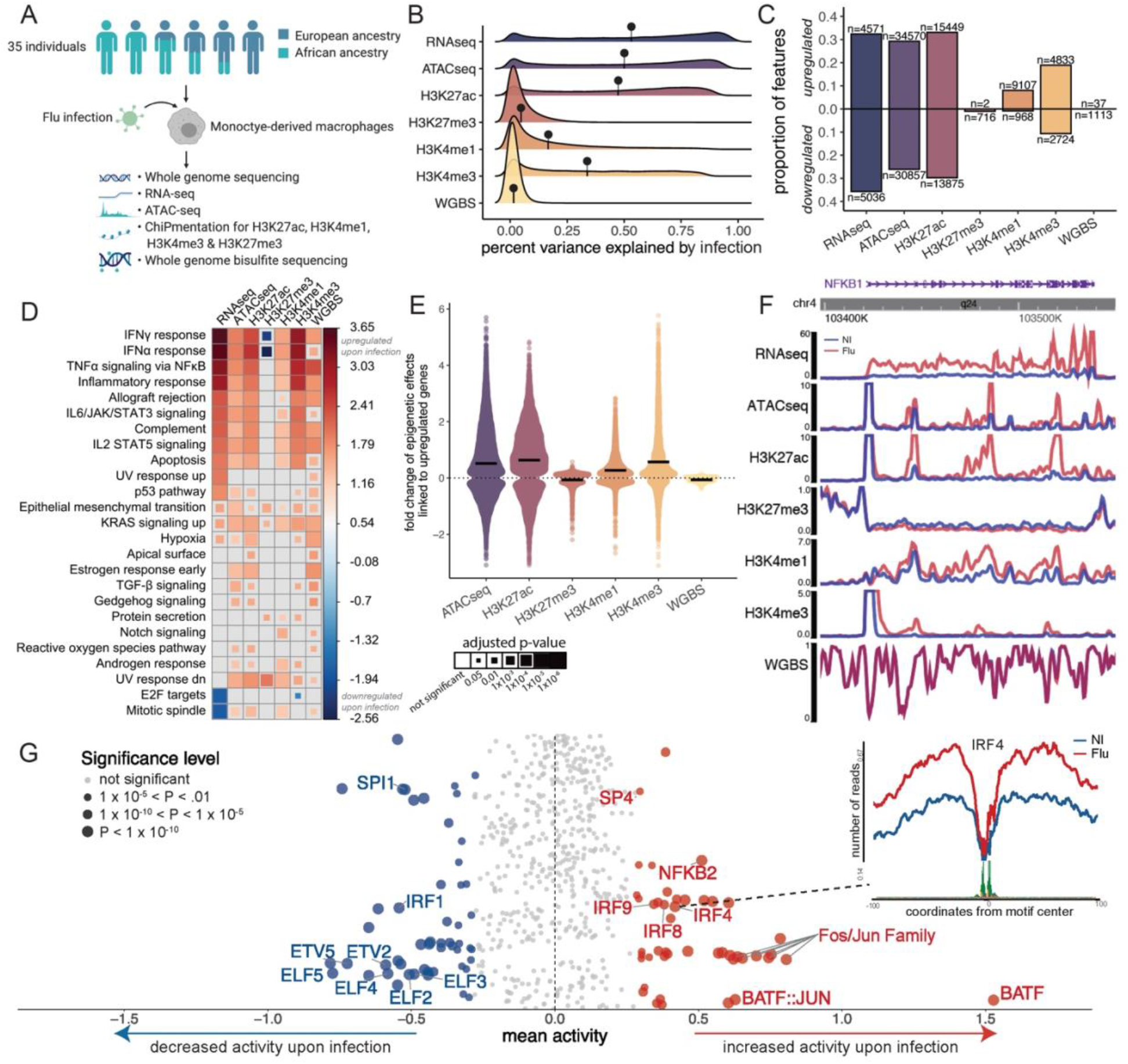
Flu infection remodels the epigenetic landscape of human macrophages. (A) Study design schematic. Monocyte-derived macrophages from 35 individuals were exposed to influenza A virus or media, for 24 hours. DNA collection and libraries for 7 types of regulatory marks were prepared and sequenced. Figure was created using BioRender.com (B) The distribution of the percent variance explained by infection for each feature in each data type. The mean is represented by the black lollipop. (C) Proportion and number of features significantly upregulated and downregulated in response to flu infection (FDR<.10, beta = ± 0.1 for WGBS and ± 0.5 for all other data types). (D) Hallmark pathways enriched among genes nearby epigenetic features (or the actual gene in case of gene expression) that respond to flu infection. Pathways shown have Benjamini-Hochberg adjusted *P* < 0.001 in at least 1 data type and a |normalized enrichment score| > 1.5 in at least 2 data types. Blue marks upregulation for that data type and red downregulation. (E) Distribution depicting the relationship between gene expression changes and epigenetic changes in response to flu infection. Mean is represented by the black line. Upregulated genes are defined as genes with beta > 0.5 and FDR<.01. Epigenetic changes are those with FDR<.01, with the exception of methylation changes for which we use a less stringent threshold (FDR<.20) due to the relatively smaller number of changes. A similar plot for downregulated genes can be found in Fig S1B. (F) The region surrounding *NFKB1*, an example of a region where gene expression and epigenetic changes occur in a coordinated fashion. (G) Transcription factor activity changes after flu infection. Upper right plot shows an example of a footprint centered on theIRF4 motif. The footprint is stronger in the flu-infected condition indicating higher levels of IRF4 activity after flu infection. Mean activity across the samples (x-axis) is plotted in the main plot. The size of the dots reflects significance levels.

We first investigated the impact of flu infection across the different data types. For DNA methylation, because most CpG sites are fully methylated and static (Pacis et al. 2015), we focused uniquely on CpG sites overlapping putative regulatory elements as identified by the chromatin segmentation program ChromHMM (Ernst and Kellis 2012) (∼7.3 million CpG sites out of a total of 19.5 million surveyed across the genome were used in all downstream analyses). Principal component analysis (PCA) on the matrices of gene expression and peak intensities revealed a strong infection effect, with NI and flu samples consistently separating on either PC1 or PC2 for most datasets (Fig. S1A). Such separation was not observed for DNA methylation or H3K27me3. To quantify the impact of flu infection on each of the molecular traits, we calculated the percent of variance explained (PVE) by infection for each feature in each data type. PVE by flu infection was highest for gene expression, chromatin accessibility, and H3K27ac histone modifications (average PVE ranging from 53-47%), followed by changes in H3K4me3 (34%) and H3K4me1 levels (17%). The least dynamic response to flu was observed for repressive marks, H3K27me3 and DNA methylation, with average PVE values across all features tested of only 5% and 2%, respectively (Fig. 1B). Consistent with the PVE analyses, 68% of all genes tested (n=9,607) were found to be differently expressed in response to flu infection (FDR<0.10 with fold change >|0.5|), Fig. 1C, Table S2). At the epigenetic level, 55% (n=65,427) and 63% (n=29,324) of regions tested changed chromatin accessibility and H3K27ac levels, respectively, in contrast to less than 0.02% for methylation and 1% of H3K27me3 levels (FDR<0.10 with fold change >|0.5|) for all data types or >|0.1| for WGBS). We also see a bias in the direction of the infection effects across the data types: repressive marks (H3K27me3 and DNA methylation) tend to be downregulated in response to flu infection (>97% of all significant changes are associated with H3K27me3 and DNA methylation losses) whereas marks associated with active enhancer and promoter regions (H3K4me1 and H3K4me3) are primarily upregulated (64-90%).

Consistent with previous work (Killip, Fodor, and Randall 2015; Ciancanelli et al. 2016), we found that genes upregulated in response to flu infection are strongly enriched for gene sets involved in interferon α and γ responses as well as the activation of inflammatory responses (normalized enrichment score (NES)>3; FDR<1×10^-6^; Fig. 1D). Our data shows that epigenetic changes in response to flu converge to the same pathways, indicating that transcriptional and epigenetic changes act in a coordinated manner to establish the immune regulatory networks required for the host response to flu infection. To further investigate the relationship between gene expression changes and epigenetic changes in response to flu infection, we asked how epigenetic features nearby genes that are up- or downregulated in response to infection respond. Regulatory elements nearby upregulated genes (Fig. 1E) show, on average, increased opening of chromatin, increased activation marks in enhancer and promoters, and a reduction of repressive marks (*P*<7.147×10^-6^ for H3K27me3; *P*<2.2×10^-16^ for all other data types; Fig. 1F for an example at the *NFKB1* locus). In contrast, regulatory elements near downregulated genes tend to be associated with closing of chromatin and the loss of activation marks (Fig S1B).

To investigate the role played by transcription factors (TF) to the epigenetic changes identified in response to IAV infection we used TF footprinting to compute TF activity scores (Fig. 1G, Table S3). TF footprinting characterizes regions where TFs are likely bound based on chromatin accessibility patterns at known TF motifs. We were particularly interest in TFs which activity levels change between NI and flu-infected samples (Fig. 1G inlet). We find that many immune-related TFs, such as those in the Fos/Jun family and Interferon Regulatory Factors (e.g., IRF4, IRF8 and IRF9), significantly increase activity after infection (*P*<1×10^-5^). Of note, we find that several ETS family members are downregulated in response to flu infection, which is concordant with ETV7 acting as a negative regulator of the type I IFN response (Froggatt et al. 2021; Pezzè et al. 2021). Unexpectedly, BATF, which is not a classical immune-related TF, showed the greatest increase in activity upon infection. Our results thus suggest that BATF likely plays a previously unappreciated role in the macrophage response to flu infection, paralleling its already established role in the induction of effector programs and epigenetic landscape of CD8 ^+^ T cells and innate lymphoid cells infected with flu (Wu et al. 2022; Lee et al. 2021; Scott-Browne et al. 2016). Collectively, our results show that transcriptional and epigenetic changes in response to flu infection are highly coordinated and likely driven by the activation of infection-induced TFs involved in the regulation of antiviral responses.

### Ancestry-associated differences across transcriptional and epigenetic responses to influenza infection

We next investigated ancestry-associated differences across the data types. To do so, we used the genotype data to estimate genome-wide levels of European and African ancestry in each sample using ADMIXTURE (v1.3.0) (Alexander, Novembre, and Lange 2009). Consistent with previous reports (Tishkoff et al. 2009), we found that self-identified African American (AF) individuals have a high proportion of European ancestry (mean = 28%, range 13%–57%; Fig. S2A). In contrast, self-identified European Americans (EU) showed virtually undetectable levels of African admixture (mean = 0.05%, range 0.001%–0.69%; Figure S2A). In all downstream analyses, African ancestry level was used as a continuous variable unless otherwise noted.

We first identified genes/regions where gene expression, accessibility, histone changes, or methylation are correlated with quantitative genetic ancestry estimates at baseline, after flu infection, or both. We termed these genes/regions as “population differentially expressed” (popDE). Combining both FDR and a multivariate adaptive shrinkage method (mash) (Storey et al. 2019; Urbut et al. 2019), we identified both shared and condition-specific popDE features for each data type (conservatively defined as genes/regions significant at a FDR<10% & local false sign rate (lfsr) < 10%) (Fig. 2A, Table S4). Mash leverages the correlation structure across conditions increasing statistical power and enabling the detection of shared popDE effects. We found that gene expression and H3K4me1 levels show the largest proportion of significant differences between ancestry groups were the most divergent between ancestry groups – 23% of genes and 21% of H3K4me1 peaks tested were classified as popDE across infected and non-infected macrophages. In contrast, only 1% of promoter-associated H3K4me3 peaks were classified as popDE at the same significance thresholds, suggesting that ancestry-associated differences in gene expression are primarily driven by variation in enhancer activity as opposed to variation at the level of core promoters. As with flu infection effects, we found that popDE effects at the transcriptional and epigenetic level were highly coordinated: genes more highly expressed in individuals with a greater proportion of African ancestry were linked to epigenetic changes indicative of increased transcriptional activity in African-as compared to European-ancestry individuals, including increased chromatin accessibility, histone acetylation levels (H3K27ac), as well as mono-and tri-methylation of H3K4 (Fig. 2B, Fig. S2B).

**Figure 2.**
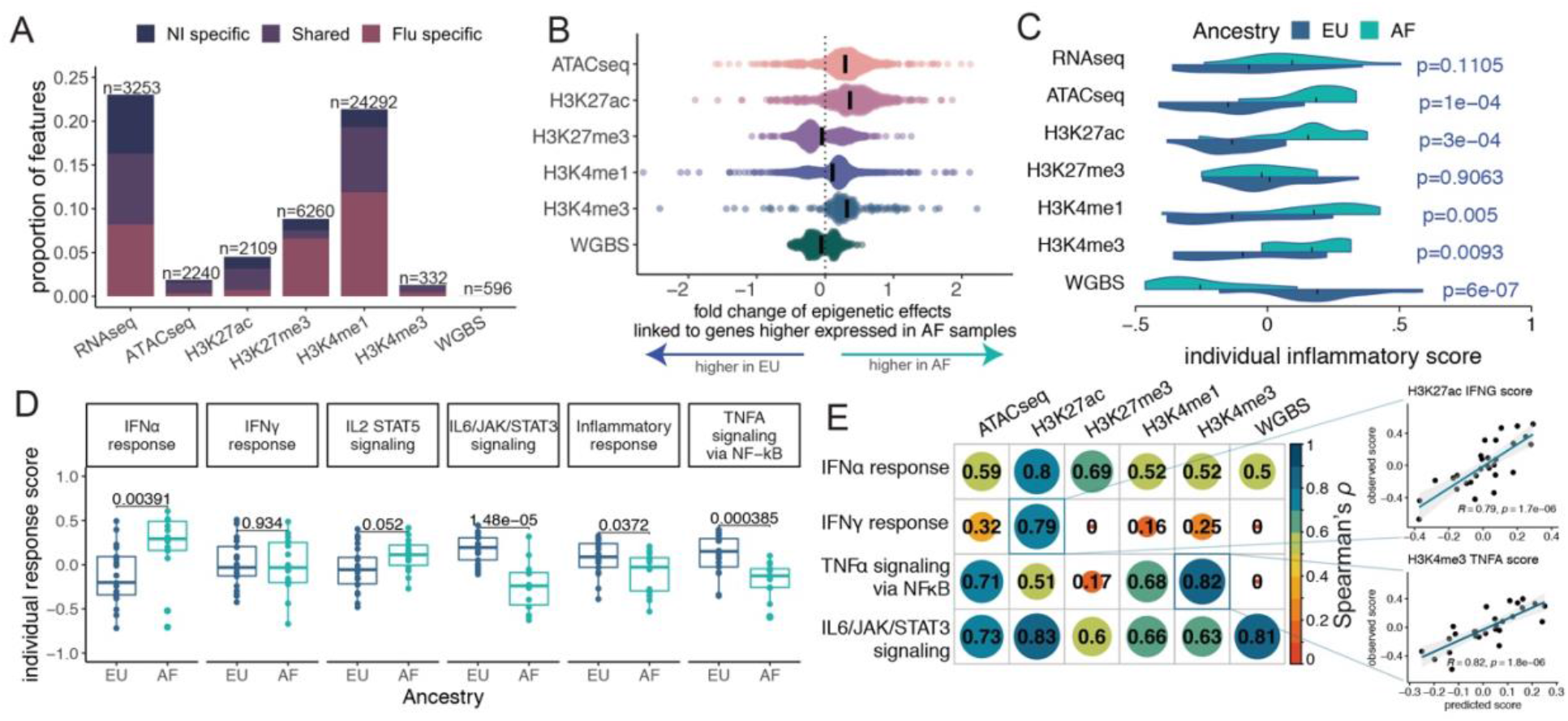
Ancestry-associated differences in the gene regulatory response to flu infection. (A) Proportion and number of popDE features that are either condition-specific (FDR<.10 and lfsr<.10 in only one condition) or shared (FDR<.10 and lfsr<.10 in both conditions). (B) Distribution depicting the relationship between popDE genes and popDE epigenetic changes. Genes more highly expressed in individuals with high proportions of African ancestry (fold change> 0.5, FDR< 0.10) are nearby popDE epigenetic regions showing increased levels of chromatin accessibility, H3K27ac, H3K4me1 and H3K4me3 in individuals with increased African ancestry. Black lines represent means. (C) Distributions of individual mean score for the Hallmark “inflammatory pathway”. A higher score value indicates a stronger expression of genes or epigenetic marks nearby genes within this inflammatory response pathway. *P* values were calculated using a Wilcoxon rank sum test. (D) Boxplots of individual transcriptional response scores for 6 immune response pathways. Pathway response levels were measured as the difference in the per individual pathways’ score between the flu-infected and non-infected conditions. (E) Pearson’s correlation of observed and predicted transcriptional response scores.

Inflammation levels have consistently been shown to vary between individuals of European and African ancestry, with an overall tendency for higher inflammation in individuals with increased African ancestry (Nédélec et al. 2016; Pennington et al. 2009; Quach et al. 2016). To evaluate if differences in inflammation could result from baseline epigenetic differences between ancestry groups, we computed a per-sample score of inflammatory activity, the “inflammation score”, which provides an estimate of the average expression or peak height of all features (genes or peaks nearby genes) in the Hallmark inflammatory response pathway (Liberzon et al. 2015). Consistent with previous reports, we found a clear trend towards higher “inflammation score” at the gene expression level in African ancestry individuals relative to Europeans in the non-infected condition (1.3-fold, albeit non-significant, *P*=0.11, Fig 2C). More strikingly, we also found an epigenetic signature of higher inflammation in individuals of African-ancestry, relative to European-ancestry individuals. Specifically, we found that increased levels of African-ancestry were strongly associated with increased levels of chromatin accessibility (*P*=1×10^-4^), H3K27ac (*P*=3×10^-4^), H3K4me1 (*P*=5×10^-3^), H3K4me3 (*P*=9×10^-3^) as well as lower levels of CpG methylation (*P*=6×10^-7^) nearby genes involved in the regulation of inflammatory responses (Fig. 2C (NI), see Fig. 2SC for similar effects in the flu-infected condition).

Our dataset provides a unique opportunity to evaluate if baseline differences in epigenetic landscape contribute to ancestry-associated differences in transcriptional response to flu. To test such hypothesis, we started by characterizing genes for which the gene expression response to infection (i.e., individual-based fold-change) significantly correlated with genetic ancestry (hereafter referred to as population differently responsive genes; or popDR). We found 2,149 popDR genes (FDR<0.20; Table S4), reinforcing the notion that genetic ancestry has a marked impact on the transcriptional response to flu (Nédélec et al. 2016; Quach et al. 2016; Randolph et al. 2021). Focusing on this set of popDR genes and on a curated set of immune pathways known to be involved in anti-viral responses (Liberzon et al. 2015), we found that (at 24 hours post-infection) individuals with higher proportions of African ancestry show a significantly stronger IFN-α response (*P*=0.004) and weaker IL6/JAK/STAT3 (*P*=1.5×10^-5^), TNFα (*P*=3.9×10-4) and inflammatory (*P*=0.0372) responses relative to EU individuals (Fig. 2D).

To evaluate if the observed differences in gene expression responses between European and African-ancestry individuals could stem from baseline differences in epigenetic profiles, we then used elastic net regression to assess the predictive power of baseline (non-infected) epigenetic levels to the transcriptional responses of the pathways described above. We found that the response to all pathways tested could be predicted with high accuracy (*R* ≥ 0.79, *P* ≤ 2×10^-6^) by the baseline levels of at least one epigenetic mark. Across the different marks, baseline levels of H3K27ac showed the most consistent predictive value across all the pathways (*R* range: 0.51 to 0.83, *P* ≤ 5×10^-3^, Fig. 2E, Fig. S2D). These results support the idea that an individual’s gene expression changes in response to flu infection are, at least in part, driven by the epigenetic landscape of the genome surrounding the gene prior to infection.

### Single nucleotide polymorphisms and short tandem repeats independently drive differences in regulatory marks

To evaluate the contribution of genetic variation to ancestry-associated differences, we mapped genetic variants that are associated with variation in gene expression or epigenetic marks across individuals (i.e., quantitative trait loci (QTL); hereafter we will refer to the mapping of the different molecular traits as the following: gene expression (eQTL), chromatin accessibility (caQTL), H3K4me1 (K4me1QTL), H3K4me3 (K4me3QTL), H3K27ac (K27acQTL), H3K27me3 (K27me3QTL), and methylation (meQTL)). To map QTL, we used a linear regression model that accounts for population structure and principal components of the expression data, thus limiting the effect of unknown confounding factors (see methods for details). Given that our sample size is too small to robustly detect trans-acting QTL, we focused our analyses on local associations that, for simplicity, we refer to as cis-QTL, defined as variants located within a gene body/peak or in the 100 kb flanking the gene/peak of interest. For methylation levels, the window was limited to ±5kb from the CpG site being tested (Banovich et al. 2014; Huan et al. 2019). We leveraged our deep whole-genome sequencing data to obtain genetic information not only on single nucleotide polymorphisms (SNPs) but also on short tandem repeats (STRs), which constitute one of the most polymorphic and abundant types of repetitive elements in the human genome (Ellegren 2004; Gemayel et al. 2010). Across individuals, we identified approximately 7.38 million SNPs and 440,000 STRs with a minor allele frequency above 5% for SNPs and 10% for STRs, which were used for QTL mapping. We identified at least one cis-eQTL (FDR<10%) for 3,880 genes (eGenes) across one or more conditions (28% of all genes tested, Fig. 3A, Fig. S3A, S3B, S3C, Table S5). eGenes identified in our study were strongly enriched among eGenes previously reported in macrophages (Nédélec et al. 2016) (5.3-fold enrichment, *P* <1×10^-15^), attesting for the robustness of our QTL results. Among epigenetic marks, H3K4me1 was associated with the largest proportion of QTL (18.5% of all peaks tested, FDR <10%), followed by chromatin accessibility (20%) and H3K4me3 (11.6%) (Fig. 3A, Fig. 3SC). For methylation levels, we found over 43,182 CpG sites associated with at least one meQTL, but, given the large number of CpG sites tested (over 7 million), the relative number of associations was the smallest of all epigenetic marks. Strikingly, across all molecular traits tested, 5-33% of features with a cis-QTL were only identified through their association with STRs and not SNPs (Fig. 3A, Fig. S3C). We next used variance partitioning to disentangle the relative contribution of STRs and SNPs to variation in gene expression and epigenetic marks (Fig. 3B, Fig. S3D). Across molecular traits, STRs contribute, on average, to 4-10% of the *cis* heritability among genes/peaks associated with at least one QTL in infected cells, not far from the amount of variance independently explained by SNPs (9-16%). Similar results were found for non-infected cells (Fig. S3D). Thus, our findings highlight the unique contribution of STRs to the genetic architecture of human gene regulation.

**Figure 3.**
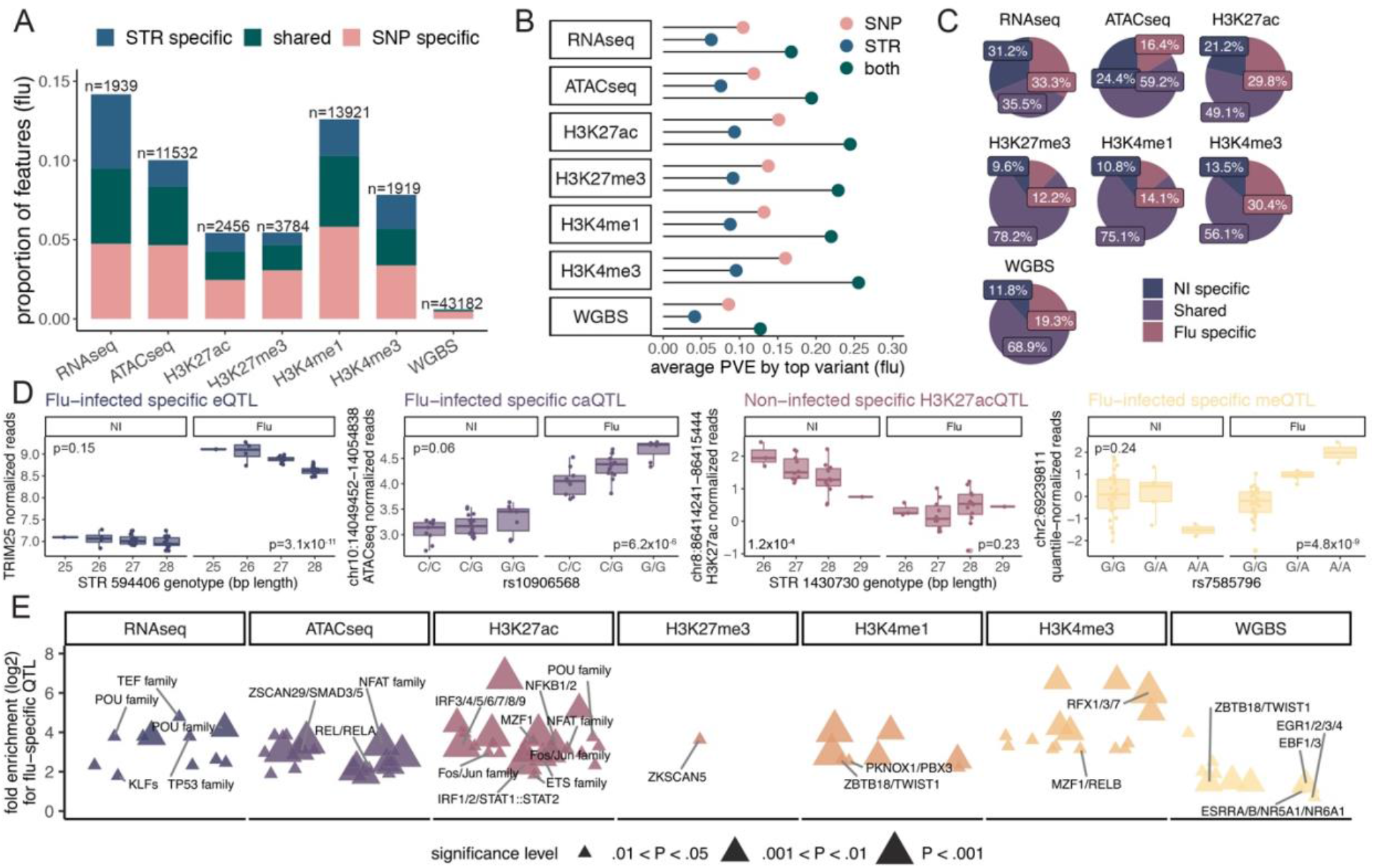
Cis-regulatory variation drives ancestry-associated differences in the transcriptional and epigenetic response to flu infection. (A) Proportion and number of genes/features associated with at least one SNP or STR QTL (in flu-infected samples, see Fig. S3C for the non-infected samples). Shared QTL were defined as those genes/features associated with a QTL at an FDR<.10 when performing the QTL mapping against SNPs and STRs separately. SNP- or STR-specific are those only identified as significant (FDR<0.1) against either SNPs or STRs (B) The mean percent variance explained by the top SNP and STR across all features in the flu-infected condition. Both is the sum of the PVE of the top SNP and top STR (C) Pie charts showing the percentage of condition specific (FDR<.10 and lfsr <.10 in only one condition with either SNP or STR) and shared QTL (FDR<.10 and lfsr <.10 in both conditions with either SNP or STR) across the data types. (D) *Far left* - An example of a flu-infected specific STR-eQTL. *Middle left* - An example of a flu-infected specific SNP-caQTL. *Middle right* - An example of a non-infected specific STR-K27acQTL. *Far right* - An example of a flu-infected specific meQTL. (E) The enrichment of TF binding sites across flu-infected specific SNP-QTL. Immune-related TF cluster names are highlighted.

We found that a large fraction of eGenes (33.3%) were only detectable in the infected condition (Fig. 3C, Fig. 3D), further reinforcing the pervasive role of gene by environment (GxE) interactions on human gene expression (Lee et al. 2014; Fairfax et al. 2014; Barreiro et al. 2012; Nédélec et al. 2016; Quach et al. 2016). Our epigenetic data expands on previous work on gene expression by showing that that GxE interactions are ubiquitous across the entire gene regulatory landscape, and not only transcription: across epigenetic marks, 12.2-30.4% of peaks (or CpG sites) showed an infection-specific QTL. We hypothesized that one potential mechanism accounting for infection-specific QTLs is that the causal SNPs/STRs disrupt the binding site of TFs that become more active in response to flu infection. To test this hypothesis, we investigated if infection-specific QTL were significantly enriched for TF footprints. We found that cis-eQTL and cis-epigenetic QTL detected only in infected cells were markedly enriched for a diverse array of immune-activating TF footprints (e.g., IRF and NfK-B family members; Fig 3E, Table S3), suggesting that many infection-specific QTL are likely to be driven by the differential binding of infection-induced TFs.

### QTL are shared across regulatory marks

To further investigate the connection between genetically regulated variation in epigenetic marks and gene expression levels, we tested if SNPs that are QTL in one data type are also QTL for the other data types. Briefly, for each condition, we took all significant SNPs (FDR <.10, Fig. S3A) for a feature and collected their corresponding p-value for all additional features. We used a cutoff of FDR<.10 to define if the SNP is also a significant QTL in other data types. We find striking patterns of sharing across the data types. For example, in non-infected macrophages, on average, 60% of QTL identified in one data type are shared with at least one other data type (ranging from minimum 45% for meQTL and at maximum 76% for K27acQTL (Fig 4A), compared to a null expectation of only 1.8% (Fig. S4A) when permuting the data; see methods for details). We find a similar pattern for the QTL identified in the flu-infected condition (Fig. S4B, Fig. S4C, Fig. S3A).

**Figure 4.**
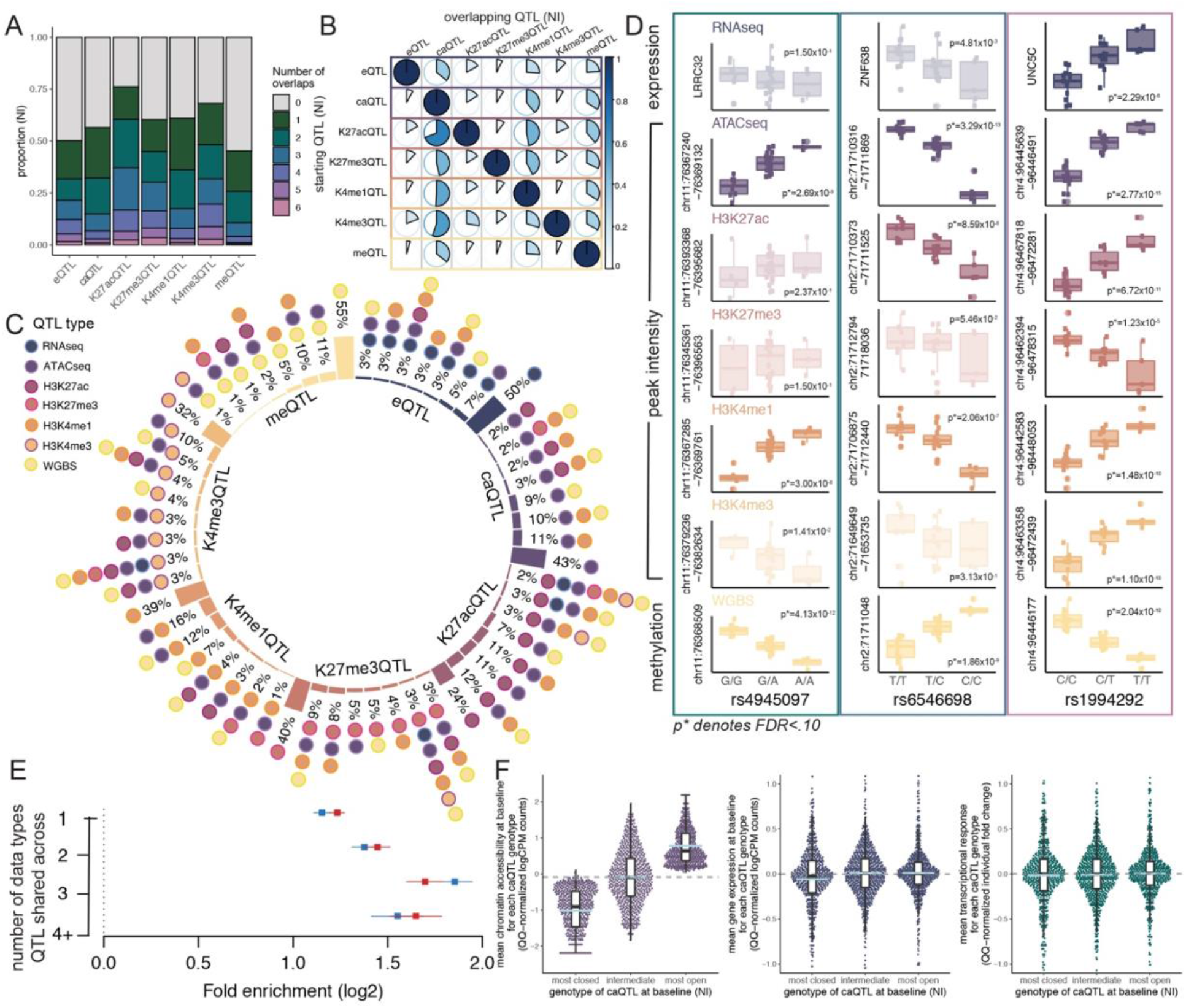
Overlap of regulatory QTLs along the cascade of gene regulatory elements. (A) The number of overlaps for each QTL type in the non-infected condition. In this figure, one (dark green) means that the QTL is only a QTL for that datatype alone. More than one overlap means that the QTL is shared with at least one other datatype, with 6 referring to cases where the QTL is shared across all datatypes. (B) The percentage of QTL in one data type that are also QTL for another data type at baseline (NI condition). The starting QTL (rows) are the QTL that are tested for sharing while the overlapping QTL (columns) are the percentage of each starting QTL that are shared with that datatype. For example, 36% of eQTL are also caQTL (row 1), while only 8% of caQTL are also eQTL (row 2). The color of each circle corresponds to the percentage of sharing. (C) The top patterns for QTL integration for each data type at baseline (NI condition). The size of the bar represents the percentage of significant QTL (FDR<.10) that share the pattern reported by the dots. (D) Examples of SNPs that are shared QTL. Grayed out plots are molecular QTL that are not significant at FDR<.10. Left: rs4945097 is a QTL for chromatin accessibility, H3K4me1 and methylation. Center: rs6546698 is a QTL for chromatin accessibility, H3K27ac, H3K4me1 and methylation. Right: rs1994292 is a QTL for all 7 data types. Notably, the T allele is associated with higher expression of *UNC5C* and epigenetic marks indicative of activated regions of the genome. (E) QTL enrichments (x axis) in actively regulated TF binding sites annotated by ATAC-seq footprinting. Error bars show 95% confidence intervals. QTLs that are shared across multiple data types are more likely to be enriched among TF footprints. (F) Association between genetically encoded baseline differences in chromatin accessibility and the magnitude of transcriptional response to IAV infection. *Left*-Meta caQTL plot at baseline condition across all caQTLs identified. For each caQTL locus, individuals were binned based on their genotype: homozygous for the genotype associated with more closed chromatin (most closed), heterozygous (intermediate), or homozygous for the allele associated with increased chromatin accessibility (most open). The light blue line marks the mean for each genotype and the gray dotted line is the median across all genotypes. We focused specifically on caQTLs nearby upregulated genes (n= 681 caQTLs associated with 506 genes) and that did not impact baseline expression levels (as shown in the middle plot). *Right*- Genotypes for chromatin accessibility levels at baseline have no impact on the transcriptional response of nearby genes.

We found consistent sharing patterns across each of the data types (Fig. 4B (non-infected), Fig. S4B (flu), Table S6). For example, in the non-infected condition, approximately 36% of eQTL are also caQTL. In fact, caQTL are the most commonly shared QTL type for all other data types, such that genetic variants impacting gene expression, a histone mark, or methylation will ∼50% of the time (range 36%-69% in the non-infected condition) also be associated with changes in chromatin accessibility (Fig. 4B, Fig. S4C). When considering genetic variants that are shared across three or more molecular traits, the most common pattern is sharing between caQTL, K4me1QTL, and meQTL (Fig. 4C, Fig. S4D), suggesting a high-level of co-regulation of chromatin accessibility, H3K4me1, and methylation levels at enhancers elements (Examples shown in Fig. 4D). Interestingly, QTL impacting multiple regulatory marks are also more likely to overlap TF footprints (Fig. 4E), supporting the idea that TFs are the primary mediators of sequence-specific regulation of gene regulatory programs (Kasowski et al. 2013; McVicker et al. 2013).

It is commonly believed that increased chromatin accessibility primes immune cells to respond faster and stronger to an immune challenge or infection (Bekkering et al. 2021; Zhang and Cao 2019), but the data supporting such a model remains circumstantial. We sought to use our genetic and epigenetic data to test this model. To do so, we focused on caQTL found in non-infected cells and their associated genes (i.e., the closest coding gene to the caQTL) and asked whether, across the genome, individuals homozygous for the genotype associated with more open chromatin showed a stronger transcriptional response as compared to individuals heterozygous or homozygous for the alleles associated with reduced opening. We limited the analyses to genetically regulated accessibility peaks associated with genes that are upregulated in response to flu infection, and that are not concomitantly eQTLs to avoid the confounding effect of baseline differences in gene expression to variation in transcriptional responses. We found that genetically driven variation in chromatin accessibility levels had no impact on the magnitude of transcriptional responses upon IAV infection (Figure 4F). This is surprising given that increased levels of open chromatin were also associated with baseline increased levels of H3K27ac and H3K4me1 (Fig. S4E), all of which have been postulated as “priming marks” for a stronger transcriptional response to infection. Overall, these data suggest that the relationship between baseline chromatin accessibility levels and transcriptional response to infectious agents is more complex than generally believed.

### Cis-regulatory variation explains ancestry associated differences to varying extents across the regulatory marks

We next sought to examine the connection between regulatory QTL and ancestry-associated differences in gene expression and epigenetic profiles. Consistent with an important role for genetics in ancestry-associated differences in the gene regulatory landscape, we found that genes/peaks associated with regulatory QTLs were more likely to be classified as popDE than expected by chance (Fig. 5A). Interestingly, the strongest enrichments were observed for epigenetic marks, especially DNA methylation, for which we observed that CpG sites with a meQTL were >28-fold more likely to be classified as popDE than those without (*P*<2×10^-16^; Fig. 5A). In contrast, the enrichment of popDE genes among genes with an eQTL, albeit significant (*P*<3.0×10^-4^), was only 1.2 fold (in NI, 1.5 fold in flu), suggesting a much greater contribution of genetics to epigenetic variation across populations compared to gene expression.

**Figure 5.**
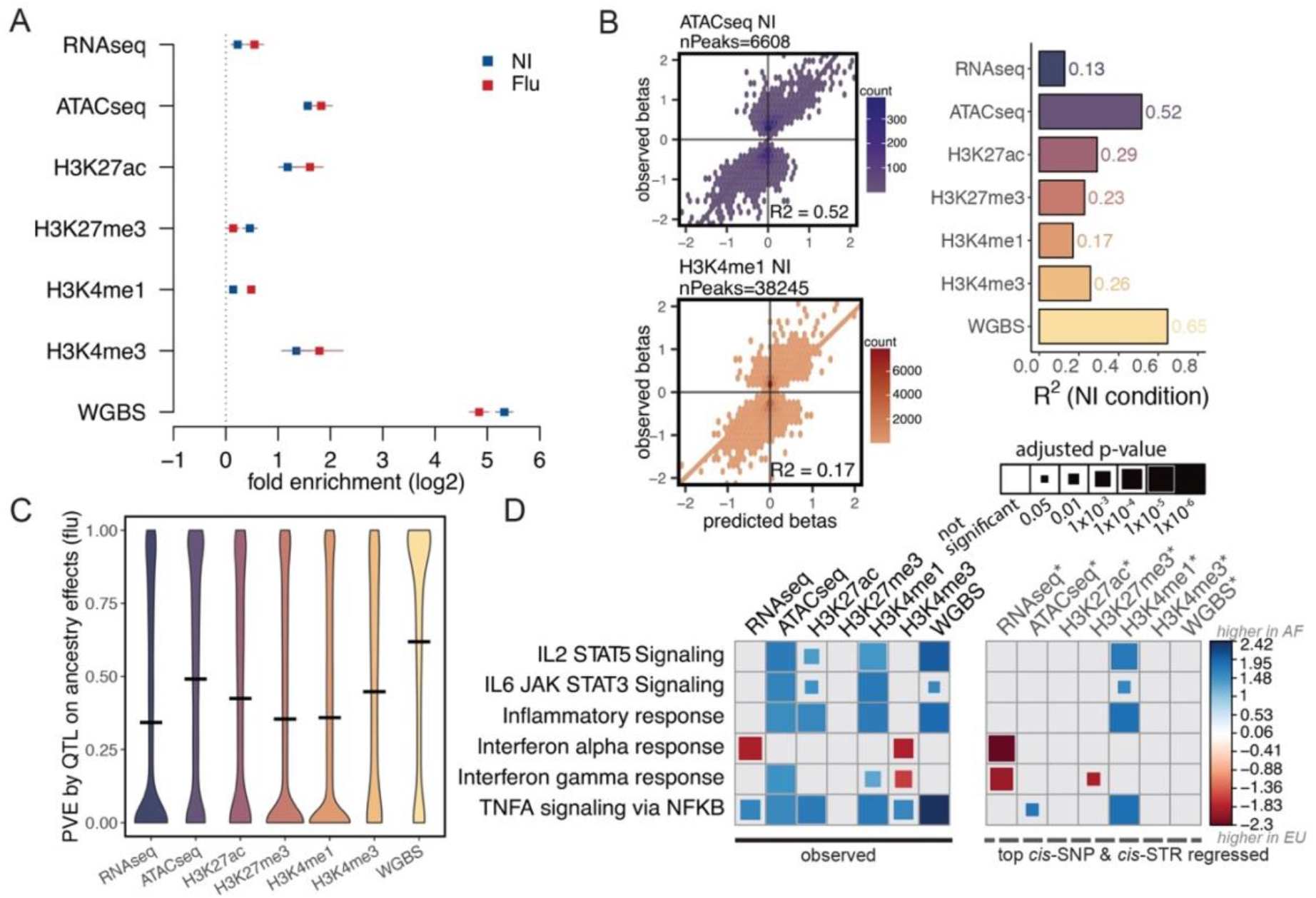
Cis-regulatory variation contributes to ancestry-associated differences. (A) The enrichment of QTL in popDE features across the data types. Log2 fold enrichments and a 95% confidence interval are plotted. (B) (Left) Examples of the correlation between the observed and predicted betas for popDE features (FDR<.10, Pearson’s correlation coefficient reported). (Right) Bar plot summarizing the correlation between observed and predicted betas for popDE features (FDR<.10) across all marks in the non-infected condition. (C) Violin plot of the percent variance explained by the top SNP- and STR-QTL on ancestry effects for each feature in each data type in the flu-infected condition. Median PVE indicated by the black line. (D) Gene set enrichment using the popDE results originally and after regressing out the top cis-SNP and cis-STR for each feature. Immune-related pathways from the Hallmark gene sets are shown. Blue indicates that the genes or features associated with genes in the pathway are more highly expressed in individuals with high levels of African ancestry. Red indicates increased expression in individuals of primarily European ancestry.

These enrichments suggest that ancestry-associated differences in gene expression are likely to be explained, at least in part, by population differences in allele frequencies at causal QTLs. To test this hypothesis, we calculated, for each of the molecular traits, the correlation between estimated and predicted genetic ancestry effects. The estimated values were obtained from our popDE analysis whereas the predicted effects were based on the effect size of the top SNP/STR for each feature and the dosage genotype for those variants across individuals (restricted to features with popDE effects). Differences in the genotype distribution between ancestry groups for the best SNP and STR explain up to 65% of the variance in genetic ancestry effect sizes across molecular traits (Fig. 5B, Fig. S5A). The strongest genetic contributions were observed for CpG site methylation and chromatin accessibility, for which the genotype of the best SNP and STR explain 65% and 52% of the variance in ancestry-associated differences in methylation and chromatin accessibility, respectively. Conversely, only 13% of the variance in gene expression differences is explained by the top SNP and STR, suggesting an important contribution of additional cis-regulatory variants, trans-regulatory variants, or environmental factors. We also calculated the change in the percent variance explained by genetic ancestry before and after regressing out the top SNP and STR. We found an analogous pattern: the lead STR and SNP plays a more significant role in explaining population differences for epigenetic marks than for gene expression (average of 62% and 49% for methylation and ATAC-seq, respectively, versus only 34% for RNA-seq, Fig. 5C, Fig. S5B (NI)).

To determine if the ancestry-associated differences in immune-related pathway activity we observed (Fig. 2C) remain significant after removing the effect of the best SNP and STR, we performed gene set enrichment analysis and compared the enrichments both before and after removing the top genetic effects (Fig. 5D). For the epigenetic effects (with the exception of H3K4me1), any baseline significant enrichment is reduced or eliminated, indicating that the top SNP and STR are important contributors to the differences in pathway activity detected between ancestry groups. For example, the observed enrichments of open chromatin and H3K27 acetylation levels near genes involved in inflammatory responses among individuals with increased African ancestry (FDR<1×10^-5^) completely disappeared (FDR>0.5) when the QTL effects were regressed out. Accounting for cis-acting genetic effects is also enough to eliminate the transcriptional differences in inflammatory response to IAV infection identified between individuals of European and African ancestry (*P*_original_ = 0.03 (Fig. 2D); P_cis-regressed_ = 0.434 (Fig S5C)). In sharp contrast, the ancestry-associated differences in type-I interferon response remain unaltered when regressing out the effects of *cis* eQTL (*P*_original_ = 0.004 (Fig. 2D); P_cis-regressed_ = 0.006 (Fig S5C)), suggesting that the ancestry-associated differences in interferon signaling are likely to be driven by environmental differences that correlate with genetic ancestry rather than by *cis* genetic variation.

### Epigenetic variants provide insight into immune-related disease risk

To evaluate the impact of regulatory QTL on susceptibility to immune-related disorders, we first assessed the colocalization (Giambartolomei et al. 2014; Wallace 2020) between regulatory QTL hits and 14 publicly available genome-wide association study (GWAS) hits for 11 immune-related diseases. For each trait, we identified the lead GWAS SNPs with p-values below 1×10^-5^ and defined a “locus” as a 100kb (5 kb window for methylation QTL) centered around the lead GWAS SNP, removing the HLA region from the analysis. We find that many epigenetic variants colocalize with variants implicated in immune-related traits (Fig. 6A, Fig. S6A, Table S7), most of which would have been missed when considering eQTL alone. Indeed, across all colocalized variants, only 7% were eQTL, the remaining corresponding to genetic variants that impact one or more epigenetic marks but not gene expression levels (e.g., Fig. 6B).

**Figure 6.**
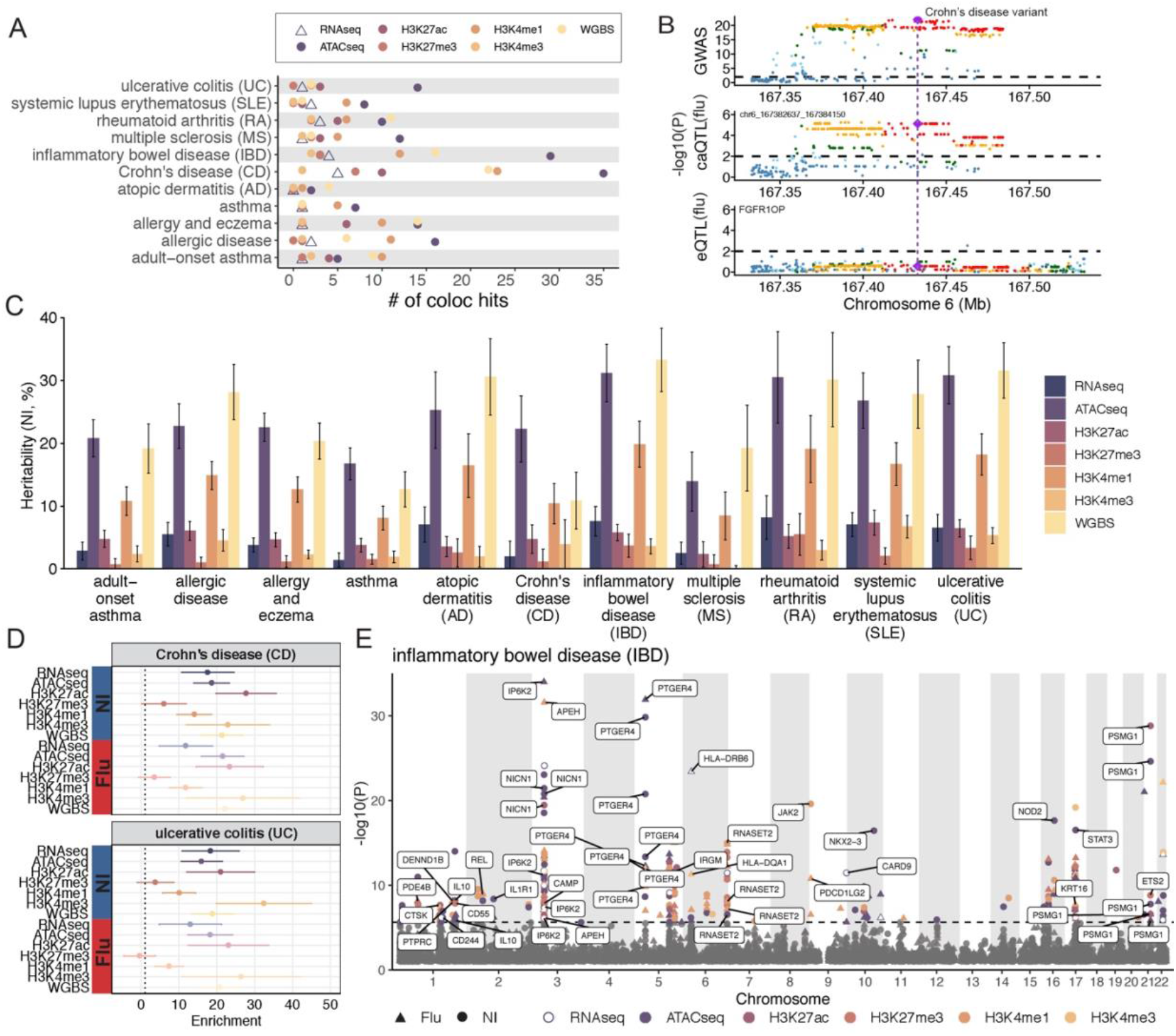
Variants controlling epigenetic marks affect immune-related disease traits. (A) Summary of colocalization results for immune related diseases. Points represent the number of significant hits defined as PP3+PP4 > 0.5 and PP4/(PP3+PP4) > 0.8 in either condition. (B) An example of a region in the flu-infected condition where a caQTL colocalizes with the GWAS variant for Crohn’s disease (the purple diamond), but the eQTL does not. *P*=0.01 is represented by the dotted line. The color of the points represents the r2, the measure of linkage disequilibrium between the SNPs. (C) Bar plots, with standard error, representing the percent of heritability explained by each of the molecular QTL in the non-infected condition. (D) Examples of heritability enrichment results. A 95% confidence interval is displayed. Enrichment results for the full 14 GWAS studies are shown in Fig. 6SD. (E) Manhattan plot showing an example of the PrediXcan results for inflammatory bowel disease susceptibility loci. Each point represents a gene or peak that is significantly associated with the disease trait. Peaks are assigned to the closest gene and labels denote genes present in the Gene Ontology immune response set (Lovering et al. 2008). The dotted line marks the *P*=0.05 Bonferroni corrected P value cutoff.

We used Stratified LD score regression (S-LDSC) (Finucane et al. 2015; Gazal et al. 2017; Bulik-Sullivan et al. 2015) to partition the heritability of complex traits and estimate heritability enrichment for each type of molecular QTL. S-LDSC is a tool for assessing how the heritability of a complex trait is partitioned among functional features, while controlling for LD, allele frequency and other baseline features. We first estimated how much heritability can be explained by each type of molecular QTL and find that chromatin accessibility and methylation QTL explain the largest percentage of heritability relative to the other data types (Fig. 6C, Fig. S6B).

We next investigated the enrichment of heritability for each molecular QTL type, estimating heritability enrichment as a ratio of the proportion of heritability explained by a particular class of regulatory QTLs divided by the proportion of SNPs that belonged to that class. We found a significant enrichment of heritability across most diseases and QTL-types tested, with the strongest enrichments observed for K27acQTL and K4me3QTL and susceptibility to Crohn’s disease and ulcerative colitis (up to 32-fold, Fig. 6D, Fig. S6C), suggesting that genetically driven epigenetic variation in macrophages plays an important role in susceptibility to gut inflammatory disorders.

Lastly, we applied S-PrediXcan to identify genes for which the component of gene expression or epigenetic values determined by an individual’s genetic profile (i.e., the regulatory QTLs identified herein) differed between cases and controls for the immune-related diseases described above (Gamazon et al. 2015). Again, we found that genetically driven differences in epigenetic marks were more frequently associated with disease status across various immune-related diseases as compared to genetically encoded variation in gene expression levels (Fig. 6E, Fig. S6D, Table S8 provides the full results of S-PrediXcan analyses across all molecular traits and 11 immune-related diseases). For IBD, for example, we found 23 genes putatively associated with disease susceptibility via changes in gene expression versus 178 genes (∼8-fold more) when focusing on genetically encoded epigenetic differences. In sum, our results consistently highlight the link between genetically encoded epigenetic variation and susceptibility to immune-related disorders.

## Discussion

Together, our results provide an extensive characterization of the gene regulatory landscape associated with variation in the immune response to flu infection between individuals of European and African ancestry. Our findings expand on previous work measuring genetic ancestry effects on the gene expression response to pathogens or immune stimuli (Nédélec et al. 2016; Quach et al. 2016; Randolph et al. 2021) by showing that many of the ancestry-associated differences in transcriptional responses to pathogens are accompanied by epigenetic differences between ancestry groups. Similar to previous findings, we found that increased levels of African ancestry are associated with a gene expression signature of increased inflammation both before and after flu infection (Brinkworth and Barreiro 2014; Pennington et al. 2009; Nédélec et al. 2016). Remarkably, we found that this signature of increased inflammatory potential among African ancestry individuals is even more accentuated when looking at the epigenetic landscape surrounding inflammation-associated genes. Other key pathways involved in the innate immune response to pathogens (e.g. Type-I interferon or TNFA signaling via NFKB) also emerged as significantly different from both an epigenetic perspective as well as in their gene expression response to flu infection between individuals of European and African ancestry. Given the central role these pathways play in the host pathogen response, our findings have potential clinical implications not only for influenza infection but also for other infectious agents.

Since samples were derived from individuals with unknown life histories and environmental exposures, the ancestry-related differences we observed could be derived from a combination of environmental and genetic factors. The integration of ancestry-effects on gene regulation with QTL analyses allowed us to demonstrate that cis-genetic variants account, in large part, for the identified ancestry-associated differences in inflammatory response. In stark contrast, ancestry-associated differences in type-I interferon response – one of the pathways most commonly identified as divergent between European and African ancestry individual (Randolph et al. 2021; Quach et al. 2016) – do not appear to be explained by differences in allele frequency of *cis* genetic variants. These data suggest, therefore, that variation in interferon responses is likely environmentally driven or explained by *trans* genetic variants that we are unpowered to identify in this study. More generally, we show that genetics contributes more to epigenetic variation at the population level than it does towards variation in gene expression, corroborating previous findings only focused on population variation in DNA methylation levels (Carja et al. 2017; Husquin et al. 2018). We speculate that this finding reflects a more direct causal role of variation in TF binding to epigenetic variation versus gene expression levels that often require the combined action of several transcriptional regulators and regulatory elements. In general, our data points to a driving role for differential TF binding in many of the molecular QTL identified (especially the epigenetic QTL), suggesting that additional effort should be invested to developing large scale datasets of TF-binding QTL, which as of now remain scarce and limited to very few TFs (Kasowski et al. 2013; Ding et al. 2014; Tehranchi et al. 2016).

Our data raises questions about the commonly accepted notion that increased chromatin accessibility at baseline allows for a stronger transcriptional response to infection (Bekkering et al. 2021; Zhang and Cao 2021). Although we cannot exclude the possibility that this is true at a limited number of loci (Alasoo et al. 2018), we show that this is not a generalizable feature across the genome. We show that an increase in chromatin accessibility prior to infection –coupled with higher levels of other activation marks, such as H3K4me1 and H3K27ac – is not in itself sufficient to “prime” cells to respond differently to a pathogenic attack. It is therefore likely that enhancer priming requires, in addition to epigenetic modification, active changes in the baseline activity of particular TFs as well as changes to the metabolic state of macrophages (Fanucchi et al. 2021). Our conclusion, however, has to be considered within the limitations of our experiment; notably the fact that we have only measured transcriptional responses at a single time point (24 hours post infection) and that we are limited in our ability to link specific enhancers to the genes that they regulate.

Finally, our results indicate that epigenetic QTL are a powerful means to identify the mechanisms of disease-associated genetic variation. About 90% of GWAS variants map to non-coding regions of the genome, suggesting that they likely affect traits through gene regulation (Hindorff et al. 2009). Despite immense efforts to characterize eQTL across thousands of individuals, tissue types and experimental conditions (GTEx Consortium 2017; Võsa et al. 2021), only ∼40% of GWAS variants colocalize with eQTLs (GTEx Consortium 2020). The modest overlap between GWAS loci and eQTLs is often attributed to the fact that many of the GWAS variants may only have an impact on gene expression during development, in specific cell types, or under environmental/experimental conditions not yet profiled. Our data indicates that epigenetic QTLs help fill this gap, by providing a means to markedly increase the number that colocalize (by about 10-fold) between GWAS variants and regulatory variants beyond those identified using eQTLs alone. We caution interpreting epigenetic variation as the causal mechanism behind variation in disease traits. We speculate, instead, that these epigenetic QTLs allow for the identification of sites associated with variation in TF binding. Therefore, they may serve as a proxy for genetic variation that, under particular environmental conditions, will have an impact on gene expression levels (Figure S7E for a schematic model). Collectively, our data indicates that our understanding of disease etiology, genetic heritability, and disease risk can be greatly increased by considering molecular traits beyond gene expression.

## STAR METHODS

Detailed methods are provided in the online version of this paper and include the following:

## SUPPLEMENTAL INFORMATION

Supplementary Information includes 6 figures and 8 tables.

## ACKNOWLEDGEMENTS

We thank Silvia Vidal from McGill University for a gift of the Influenza strain. We thank all members from the Barreiro and Bourque labs for their comments on the paper. This work was supported by National Institute of Health Research grants R01-GM134376 and P30-DK042086 to L.B.B. It is also supported by a Canada Institute of Health Research (CIHR) program grant (CEE-151618) for the McGill Epigenomics Mapping Center, which is part of the Canadian Epigenetics, Environment and Health Research Consortium (CEEHRC) Network, to G.B., L.B.B. and T.M.P. K.A.A. is supported by a grant to University of Chicago from the Howard Hughes Medical Institute through the James H. Gilliam Fellowships for Advanced Study program. G.B. is supported by a Canada Research Chair Tier 1 award, a FRQ-S, Distinguished Research Scholar award and by the World Premier International Research Center Initiative (WPI), NEXT, Japan. The Canadian Center for Computational Genomics (C3G) is supported by a Genome Canada Genome Technology Platform grant. Computational resources were provided by the University of Chicago Research Computing Center (Barreiro team) and Calcul Québec and Compute Canada (Bourque team). Figures 1A and 7E were created with BioRender.com.

## AUTHOR CONTRIBUTIONS

L.B.B, G.B and T.M.P conceived the project. L.B.B. directed the study. V.Y., R.S., A.P (Albena), and M.S. performed experimental work. K.A.A. led the computational analyses, with contributions from Y.L, A.P (Alain), S.G., Z.M., K.L, C.G., X.C., X.H. and Y.L. A.P. (Alain) and D.L. developed and implemented the EpiVar browser with help from R.G., D.B., and D.B. K.A.A. and L.B.B. wrote the manuscript, with input from all authors.

## STAR METHODS

## KEY RESOURCES TABLE

TBD

## RESOURCE AVAILABILITY

### Lead contact

Reagent and resource requests should be addressed and will be fulfilled by the Lead Contacts, Luis Barreiro and Guillaume Bourque.

### Materials availability

This study did not generate new unique reagents.

### Data and code availability

Sequence data has been deposited at the European Genome-phenome Archive (EGA), under accession numbers EGAD00001008422 (RNA-seq, ATAC-seq and ChIPmentation) and EGAD00001008359 (WGS and WGBS). We also constructed a versatile QTL browser (https://computationalgenomics.ca/tools/epivar), which allows users to explore and visualize mapped QTLs for gene expression, chromatin accessibility, histone modifications and DNA methylation.

All original code is currently available at https://github.com/katiearacena/EU_AF_ancestry_flu_code and will be deposited at Zenodo by the date of publication.

Any additional information required to reanalyze the data reported in this paper is available from the lead contact upon request.

## EXPERIMENTAL MODEL AND SUBJECT DETAILS

### Sample collection

Buffy coats from 39 healthy donors were obtained from the Indiana Blood Center (Indianapolis, IN, USA). A signed written consent was obtained from each participant and the project was approved by the ethics committee at the CHU Sainte-Justine (protocol #4022). All individuals recruited in this study were males, self-identified as African-American (AF) (n = 19) or European-American (EU) (n = 20) between the age of 18 and 54 years old. The average age across AF and EU samples was similar (38.7 years for AF versus 38.6 years for EU). We only collected male samples to avoid the potentially confounding effects of sex-specific differences in immune responses to infection. Only individuals self-reported as currently healthy and not under medication were included in the study. In addition, each donor’s blood was tested for Hepatitis B, Hepatitis C, Human Immunodeficiency Virus (HIV), and West Nile Virus, and only samples negative for all of the tested pathogens were used.

### Monocytes isolation and macrophages generation

Blood mononuclear cells were isolated by Ficoll-Paque centrifugation. Monocytes were purified from peripheral blood mononuclear cells (PBMC) by positive selection with magnetic CD14 MicroBeads (Miltenyi Biotech) using the autoMACS Pro Separator. The purity of the isolated monocytes was verified using an antibody against CD14 (BD Biosciences) and only samples showing > 90% purity were used to differentiate into macrophages. To generate the monocytes-derived macrophages (MDM), the cells were cultured for 6 days in RPMI-1640 (Fisher) supplemented with 10% heat-inactivated FBS (FBS premium, US origin, Wisent), L-glutamine (Fisher), gentamicin (10ug/mL LifeTechologies) and M-CSF (20ng/mL; R&D systems) and incubated at 37°C and 5% CO_2_. Cell cultures were fed every 2 days with complete medium.

### Infection of macrophages

On day 6, the macrophages were harvested with CellStripper (Corning), counted, replated with the fresh media (previously mentioned) without antibiotic and incubated overnight. The next day, the cells were infected at a multiplicity of infection (MOI) of 0.1 for Influenza A virus strain *PR8WT* (Flu). A control group of non-infected macrophages (NI) was treated the same way but with only medium without virus. For some samples, we added Mock at the same volume as for the Flu and NI conditions. 24 hr post-infection, the cells were collected for downstream experiments.

### gDNA extraction

Genomic DNA extraction was performed on 0.6 to 7 million (from NI or Flu macrophages) using the DNeasy Blood & Tissue kit (Qiagen). The genomic DNA was quantified using Quant-iT PicoGreen ds DNA Assay Kit (ThermoFisher Scientific).

### Whole genome sequencing (WGS)

Libraries were generated from 400 ng of genomic DNA fragmented to 300–400 bp peak sizes using the Covaris focused-ultrasonicator E210. Library preparation was done using NxSeq AmpFREE Low DNA Library Kit (Lucigen) according to the manufacturer’s instructions. The libraries were size selected using Ampure XP Beads (Beckman Coulter) and quantified using the KAPA Library Quantification kit - Universal (KAPA Biosystems). Sequencing of the WGS libraries was performed on the Illumina HiSeqX system using 150-bp paired-end sequencing.

### Whole genome bisulfite sequencing (WGBS)

Libraries were generated from 1500 ng of genomic DNA spiked with 0.1% (w/w) unmethylated λ DNA (Roche Diagnostics) fragmented to 300–400 bp peak sizes using the Covaris focused-ultrasonicator E210. Library preparation was done using NxSeq AmpFREE Low DNA Library Kit (Lucigen) according to manufacturer’s instructions, followed by bisulfite conversion with the EZ-DNA Methylation Gold Kit (Zymo Research) according to the manufacturer’s protocol. Libraries were amplified by 6 cycles of PCR using the Kapa Hifi Uracil + DNA polymerase (KAPA Biosystems) according to the manufacturer’s protocol. The amplified libraries were size selected using Ampure XP Beads (Beckm an Coulter) and quantified using the KAPA Library Quantification kit - Universal (KAPA Biosystems). Sequencing of the WGBS libraries was performed on the Illumina HiSeqX system using 150-bp paired-end sequencing.

### RNA extraction

Macrophages were directly lysed from the culture plate with 1mL of Qiazol from 0.5 to 2.5 million cells (NI, Flu and Mock) and extracted using the miRNeasy kit (QIAGEN) following the manufacturer instruction. RNA integrity was assessed with the Agilent 2100 Bioanalyzer System (Agilent Technologies).

### RNA sequencing (RNA-Seq)

RNA library preparations were carried out on 100-500 ng of RNA with RIN 1.2 to 9.8 using the Illumina TruSeq Stranded Total RNA Sample preparation kit, according to manufacturer’s protocol. The libraries were size-selected using Ampure XP Beads (Beckman Coulter) and quantified using the KAPA Library Quantification kit - Universal (KAPA Biosystems). Sequencing of the RNA-Seq libraries was performed on the Illumina NovaSeq 6000 system using 100-bp paired-end sequencing.

### ATAC-seq

ATAC-seq library preparation was performed using the Omni-ATAC protocol (Corces et al. 2017). 50,000 macrophages (from NI, Flu and Mock conditions) were resuspended in 1 ml of cold ATAC-seq resuspension buffer (RSB; 10 mM Tris-HCl pH 7.4, 10 mM NaCl, and 3 mM MgCl_2_ in water). Cells were centrifuged at 500 g for 5 min in a pre-chilled (4 °C) fixed-angle centrifuge. After centrifugation, supernatant was aspirated and cell pellets were then resuspended in 50 μl of ATAC-seq RSB containing 0.1% IGEPAL, 0.1% Tween-20, and 0.01% digitonin by pipetting up and down three times. This cell lysis reaction was incubated on ice for 3 min. After lysis, 1 ml of ATAC-seq RSB containing 0.1% Tween-20 (without IGEPAL and digitonin) was added, and the tubes were inverted to mix. Nuclei were then centrifuged for 10 min at 500 rcf in a pre-chilled (4°C) fixed-angle centrifuge. Supernatant was removed and nuclei were resuspended in 50 uL transposition mix (2x TD Buffer, 100 nM final transposase, 16.5 uL PBS, 0.5 uL 1% digitonin, 0.5 uL 10% Tween-20, 5 uL H2O). Transposition reactions were incubated at 37 °C for 30 min in a thermomixer with shaking at 1000 rpm. Reactions were cleaned up with Zymo DNA Clean and Concentrator 5 columns. Primers (i5 and i7) were added by amplification (12 cycles) using NEBNext 2x MasterMix. Sequencing of the ATAC-seq libraries was performed on the Illumina NovaSeq 6000 system using 100-bp paired-end sequencing.

### ChIPmentation

#### Crosslink step

For ChIPmentation, 1 to 5 million macrophages (from NI, Flu and Mock conditions) were washed in cold PBS prior proceed the cross-linking of DNA with formaldehyde (0.75%) by shaking the tube for 10 min at RT and adding Glycine (125nM) for additional 5 min. Cells were washed with cold PBS and centrifuged for 5 minutes at 2500 xg at 4℃. The supernatant was discarded and the cell pellet immediately frozen at −80℃.

After cell lysis, sonication of nuclei was performed on a BioRuptor UCD-300 targeting 150-500 bp size. Immunoprecipitation of the histone marks H3K27ac, H3K4me1, H3K27me3 and H3K4me3 was performed following the Auto-ChIPmentation protocol for Histones (Diagenode inc, Denville, USA) according to the manufacturer’s instructions. The libraries were size selected using Ampure XP Beads (Beckman Coulter) and quantified using the KAPA Library Quantification kit - Universal (KAPA Biosystems). Sequencing of the ChIPmentation libraries was performed on the Illumina NovaSeq 6000 system using 100-bp paired-end sequencing

## QUANTIFICATION AND STATISTICAL ANALYSIS

### WGS processing and genotyping

Raw reads were trimmed using Skewer (Jiang et al. 2014) and the resulting reads were aligned to the hg19 human reference genome using BWA-MEM (H. Li and Durbin 2009). Insertion/deletion realignment and base quality score recalibration were performed using GATK (McKenna et al. 2010) and duplicates were marked using Picard (http://broadinstitute.github.io/picard/). We used GATK’s *HaplotypeCaller* to perform SNV and INDEL calling. We filtered the joint genotyped file to exclude non-autosomal and non-biallelic variants. Additionally, we removed SNPs that had a call rate of <90% across all samples, that deviated from Hardy–Weinberg equilibrium at p < 10^-5^, and with minor allele frequency less than 5%. We used the resulting 7,383,243 SNPs in QTL mapping and other downstream analyses. We annotated the SNPs using dbSNP (human_9606_b151_GRCH37p13) (Sherry ST, Ward MH, Kholodov M, Baker J, Phan L, Smigielski EM 2001).

### Estimation of genome-wide admixture levels

We used the clustering algorithm ADMIXTURE (v1.3.0) to calculate the percentage of African and European ancestry in each individual (Alexander, Novembre, and Lange 2009). Notably, we only obtained genotyping data for 17/19 self-identified African individuals and 18/20 European individuals, thus, we only calculated admixture estimates for samples we had data (n=35). We included Yoruba (YRI) and European (CEU) individuals from the 1000 Genomes reference panel and estimated ancestry proportions using K=2 ancestral clusters. We applied Genotype Harmonizer (Deelen et al. 2014) to align and combine the 1000 genomes reference data. We used 362,075 unlinked SNPs (r^2^ between all pairs < 0.1) to estimate genetic ancestry. ADMIXTURE analyses showed that 3 AF individuals were likely mislabeled by the blood center as they presented 99.9% of European ancestry. Ancestry labels were adjusted accordingly resulting in 14 African American and 21 European American individuals. Estimated ancestry proportions for each individual were used to calculate population differences unless specified otherwise.

### RNA-seq data processing

Adaptor sequences and low-quality score bases (Phred score < 30) were first trimmed using Trimmomatic (Bolger, Lohse, and Usadel 2014). The resulting reads were aligned to the hg19 human reference genome assembly, using STAR (Dobin et al. 2013). Read counts are obtained using HTSeq (Anders, Pyl, and Huber 2015) with parameters -m intersection-nonempty - stranded=yes.

### ChIPmentation and ATAC-seq data processing

ChIPmentation and ATAC-seq reads were first trimmed for adapter sequences and low-quality score bases using Trimmomatic (Bolger, Lohse, and Usadel 2014). The resulting reads were mapped to the human reference genome (hg19) using BWA-MEM (H. Li and Durbin 2009) in paired-end mode at default parameters. Only reads that had a unique alignment (mapping quality > 20) were retained and PCR duplicates were marked using Picard tools (https://broadinstitute.github.io/picard/). Peaks were called using MACS2 software suite (Y. Zhang et al. 2008).

### Filtering phenotype data

In our RNAseq dataset, we excluded any genes that did not have an average RPKM > 2 in Flu *or* non-infected samples. For the CHIPseq and ATACseq datasets, we calculated median peak size and required 50% of median value overlap for peaks to be called as the same peak between samples using bedtools merge (Quinlan and Hall 2010). We then filtered to exclude peaks that were not present in ≥ 50% of Flu *or* non-infected samples, and those that fall within blacklisted regions (Amemiya, Kundaje, and Boyle 2019). The number of features remaining after these thresholds are present in the table below.

| Data | Median peak size | 50% of median for peak merging | Total # of samples included in analysis | $\geq 50\%$ NI condition threshold | $\geq 50\%$ Flu condition threshold | Features *BEFORE* filtering | Features *AFTER* filtering |
| --- | --- | --- | --- | --- | --- | --- | --- |
| RNAseq | N/A | N/A | 70 | N/A | N/A | 57905 | **14122** |
| ATACseq | 608 | 304 | 70 | 18/35 | 18/35 | 974189 | **118201** |
| H3K27ac | 349 | 174.5 | 58 | 15/29 | 15/29 | 608338 | **46657** |
| H3K27me3 | 592 | 296 | 56 | 14/28 | 14/28 | 694827 | **70675** |
| H3K4me1 | 498 | 249 | 60 | 15/30 | 15/30 | 749325 | **113584** |
| H3K4me3 | 843 | 421.5 | 54 | 14/27 | 14/27 | 144765 | **25568** |

We used featureCounts to calculate the number of reads for each genomic feature for each sample (Liao, Smyth, and Shi 2014). We used the resulting counts matrices for all downstream analyses.

### Partitioning the genome using ChromHMM

We generated genome-wide, gene regulatory annotation maps for noninfected and flu infected MDMs using the ChromHMM chromatin segmentation program (Ernst and Kellis 2012). We used samples for which there was data for all 4 histone marks (n= 27 samples, 10 AF, 17 EU) and 7 emission states. We used ChromHMM profiles from the Roadmap Epigenetics project to annotate our results (Roadmap Epigenomics Consortium et al. 2015).

### Whole genome bisulfite sequencing data processing

Adaptor sequences and low-quality score bases were first trimmed using Trimmomatic (Bolger, Lohse, and Usadel 2014). The resulting reads were mapped to the human reference genome (hg19) and lambda phage genome using Bismark (Krueger and Andrews 2011), which uses a bisulfite converted reference genome for read mapping. Only reads that had a unique alignment were retained and PCR duplicates were marked using Picard tools (https://broadinstitute.github.io/picard/). Methylation levels for each CpG site were estimated by counting the number of sequenced C (‘methylated’ reads) divided by the total number of reported C and T (‘unmethylated’ reads) at the same position of the reference genome using Bismark’s methylation extractor tool. We performed a strand-independent analysis of CpG methylation where counts from the two Cs in a CpG and its reverse complement (position on the plus strand and position i+1 on the minus strand) were combined and assigned to the position of the C in the plus strand. To assess MethylC-seq bisulfite conversion rate, the frequency of unconverted cytosines (C basecalls) at lambda phage CpG reference positions was calculated from reads uniquely mapped to the lambda phage reference genome.

We obtained methylation counts for 19,492,906 loci. Due to the high coverage of the data, we opted to not perform smoothing. To reduce the total number of statistical tests performed, we limited our analyses to CpG sites in open chromatin regions using ChromHMM data (states E3 - E7). We also excluded C nucleotides that overlapped with SNPs identified in the whole genome sequencing data for our samples. After these filtering steps we analyzed methylation levels across 7,463,164 CpG sites.

### Infection effects: Infection-Related Differential Effects

We used all samples we had collected data for, not just those with genotyping data to calculate infection effects. Note that this only increased the sample size for RNAseq data (from n=35 to n=39 individuals). The sample size for all other data types remained the same. For RNAseq, ATACseq and CHIPseq datasets, we calculated normalization factors to scale the raw library sizes using calcNormFactors in edgeR (v 3.28.1) (Robinson, McCarthy, and Smyth 2010). We used the voom function in limma (v 3.42.2) to apply these factors, estimate the mean-variance relationship and convert raw read counts to logCPM values. Because samples were sequenced on different flowcells at different times (i.e., hereafter defined as “Batch”) we regressed out these putative batch effects by fitting a linear model that estimates the technical effect of sequencing batch on the different datasets. We kept the residuals from this model (i.e, batch-corrected “expression” estimates) using the residuals.MArraLM function.

To calculate global infection effects for RNAseq, ATACseq and CHIPseq datasets, batch-corrected read counts of samples corresponding to the same individual were compared in a paired design by introducing individuals as additional covariates. The following model was run using limma for each data type independently:

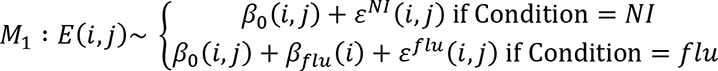

Here, *E(i,j)* represents the batch-corrected estimate of each feature *i* for individual *j* and *β_0_(i,j)* represents the intercept corresponding to feature *i* and individual *j* (i.e., the expectation of gene or peaks *i*’s expression level in the non-infected sample for individual *j*). *β_flu_(i)* is the effect of flu infection on feature *i*.

We performed 1000 permutations obtained by randomly reshuffling the condition labels in each condition in order to estimate FDR using the qvalue R package (v 2.18.0) (Storey et al. 2019).

#### Identification of differentially methylated loci

We identified differentially methylated loci (DML) in response to flu infection using the R package DSS and a fixed effects model (Park and Wu 2016). We used the DMLfit.multiFactor function in DSS, using the same model described above (M1). We performed 10 permutations and FDR correction using the same approach detailed above.

### Percent Variance Explained by Infection

The R package relaimpo (v 2.2-3) was used in order to calculate the relative contribution of each predictor in the infection effects linear models to the R^2^. (Grömping 2006). The same batch corrected counts matrices and weights were used as before with the exception of the methylation loci, which were additionally filtered to remove sites that did not have coverage ≥4 sequence reads in at least half of the non-infected or Flu-infected samples and those with 0 variation across all NI or Flu samples. DSS accounts for both low coverage and variation which is why these sites were previously included in the model. The same model (M1) was run for all datatypes.

### GSEA of Infection effects and popDE effects

The R package fgsea (Korotkevich et al. 2021) was used to perform gene set enrichment analysis (GSEA) to determine which biological pathways were enriched or depleted among DE genes/regions and popDE genes/regions. We connected CpG loci, CHIPseq peaks, and ATACseq peaks to the nearest gene using the R package ChIPseeker (Yu, Wang, and He 2015) (using the default parameters). For GSEA each gene can only be included once. Thus, in situations where more than one peak was mapped to the same gene, we kept the peak with the highest t-statistic when modeling flu-infection effects or popDE effects. For the WGBS infection effects we used the difference between CpG methylation in the flu-infected and non-infected conditions to perform GSEA since the Wald test statistic from the model does not indicate the direction of the effect. For the WGBS popDE GSEA the Wald test statistic was used. GSEA were performed against the Hallmark gene set (Liberzon et al. 2015).

### Relationship between expression and epigenetic changes in response to infection

To evaluate the relationship between gene expression changes and epigenetic changes in response to flu infection we connected peaks and CpG loci to the nearest gene using the R package ChIPseeker using the same parameters as detailed previously (Yu, Wang, and He 2015).

We first subset on upregulated genes defined as those genes with beta> 0.5 & FDR < .01. Additionally, we subset to include only epigenetic marks that change in response to flu infection using FDR < .20 for CpG loci and FDR<.01 for all other marks. We then evaluate in which direction the epigenetic features associated with genes upregulated in response to flu infection change. The same analysis is done using downregulated genes (beta < −0.5 and FDR < .01). We used a Wilcoxon test to determine significance levels using peaks for all genes (not just those upregulated and downregulated) as the null.

### Transcription Factor activity scores

Footprints were called using HINT-ATAC from the Regulatory Genomics Toolbox (Z. Li et al. 2019) on the subset of peak regions called using MACS2 (Y. Zhang et al. 2008). Footprint calling was performed by first merging aligned ATACseq reads within each condition using *samtools merge,sort,index*. A meta-footprint set was created for each pair by merging the respective footprint calls with *bedtools merge* (Quinlan and Hall 2010). Using this meta-footprint set, transcription factor motif matching was performed on the subset of regions falling within meta-footprints.

Motif matching was done using the JASPAR CORE Vertebrates set of curated position frequency matrices (Sandelin et al. 2004). Because of similarity across TF motifs, we chose to group TFs into clusters based on similarity. To do this, we first computed pairwise TOMTOM (Gupta et al. 2007) *E* value metrics to assess motif similarity. The log10 *E* values were then used as distance metrics for hierarchical clustering (base *R*; hclust(method=”ward.D2”)). A cutoff height of 10 (base *R*; cutree(h=10)) was used to define TF motif clusters, resulting in a total of 200 clusters which were used for the TF enrichment analysis described later.

Using the set of motif match regions for each TF, motif count enrichment was performed using the *rgt-motifanalysis enrichment* function. Background regions were defined as all meta-footprints. Foreground regions are the footprints overlapping regions of the genome of interest. A two-sided Fisher’s exact test was computed from the output frequencies of motif occurrences within the foreground and background regions (base *R*; fisher.test(alternative=”two.sided”)). P values were corrected using the Benjamini Hochberg method (Benjamini and Hochberg 1995).

Using the set of motif match regions for each TF, activity analysis was performed with the RGT *differential* function. The activity score metric is described further in (Z. Li et al. 2019). Parameters for footprinting, motif matching, and differential activity analysis were set as default. Activity statistics were calculated per sample between conditions using the ATACseq profiles of each sample independently. Combined activity scores were computed as the mean across samples, and meta p-values were calculated by Fisher’s combined probability test (python scipy.stats.combine_pvalues) to summarize across all samples.

### Correcting for technical effects in popDE, popDR and QTL mapping analysis

All popDE, popDR and QTL mapping analyses were performed on count matrices corrected for age and potential sequencing batch effects. Age and batch correction were done separately for NI and Flu-infected samples. We started by calculating normalization factors to scale the raw library sizes using calcNormFactors in edgeR (v 3.28.1) (Robinson, McCarthy, and Smyth 2010). Then, we used the voom function in limma (v 3.42.2) to apply these factors, estimate the mean-variance relationship and convert raw read counts to logCPM values. Batch effects, which are categorical variables, were regressed out using ComBat from the sva Bioconductor, fitting a model that also includes age (mean centered) and admixture. We subsequently regressed out age effects using limma.

### Detection of population differentially expressed (popDE) features

Using the age and batch corrected matrices described above, we used limma to detect the effect of African admixture for RNAseq, ATACseq and CHIPseq datasets using the following nested model for each data type independently:

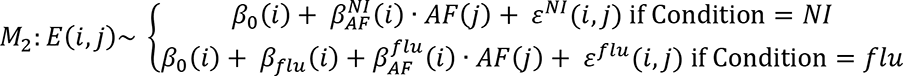

Here, *E(i,j)* represents the age and batch corrected estimate of feature *i* for individual *j*, *β_0_(i)* is the global intercept accounting for the expected expression of feature *i* in a 100% European-ancestry non-infected individual, 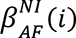 and 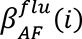 indicate the effects of African admixture on feature *i* within each condition. The model was fit using limma and the estimates of 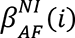 and 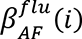 of the genetic ancestry effects were extracted across all features. We used 1000 permutations obtained by randomly reshuffling admixture estimates in order to estimate FDR, as described in detail above.

Because of the different nature of the methylation data (i.e., percentage methylation per CpG site instead of counts) we used DSS (Dispersion Shrinkage for Sequencing data) instead of limma to model population differentially methylated loci. DSS is specifically designed for the analyses of bisulfite sequencing (BS-seq) differential methylation. The core of DSS is a procedure based on Bayesian hierarchical model to estimate and shrink CpG site-specific dispersions, then conduct Wald tests for detecting differential methylation. We used the same model as M2 described above but including age and batch as covariables. We permuted the 10 times by shuffling the admixture estimates to obtain null p value distributions for FDR calculations.

To increase our power to detect condition and shared effects we applied Multivariate Adaptive Shrinkage in R (mashr v0.2.28) (Urbut et al. 2019) to the outputs of the popDE effects for each data type independently. For each condition, effect sizes were obtained from limma and standard error of the effect size was calculated by multiplying the square root of the posterior variance (s2.post) of each feature by the unscaled standard deviation for the effect size of interest for that feature (stdev.unscaled). NI and Flu effect sizes and standard error of effect sizes were formatted into n x m matrices, where: n = number of features for each data type, m = 2 conditions (NI and Flu).

We estimated the null correlation of the data using the “estimate_null_correlation_function” in mashr. We included canonical covariance and data-driven covariance matrices. The data-driven covariance matrix is the top 5 PCs from a PCA performed on the significant (local false sign rate (lfsr) < .05) signals identified in the condition-by-condition model learned from our data in the mash model fit. We then fit the mash model using the mash function. For the methylation data we made some modifications to the mash procedure. First, we removed any NAs from the DSS results, resulting from insufficient coverage at a particular CpG site. As described above, DSS uses a Wald statistical test to test each gene/CpG site for differential methylation, so we used the Wald test statistic as the effects, setting all standard errors to 1. Instead of using all the tests to estimate the null correlation structure like the other datasets, we obtained a random subset of 200,000 tests and applied the estimate_null_correlation_simple function. As with the other data types, we included canonical covariance and data-driven covariance matrices and fit the mash model on all the tests performed.

After running mash, we conservatively used both qvalue and local false sign rate (lfsr) to determine if popDE effects were condition-specific (i.e., only showing an effect in the non-infected or flu-infected conditions) or shared (i.e., showing an effect in both conditions). Specifically, we require popDE features to have both FDR <.10 and lfsr < .10 in only one condition to be considered condition specific. popDE features are shared if FDR < .10 and lfsr <.10 in both conditions.

### Detection of population differentially responsive features

We used the age and batch corrected count matrices and weights to model the effects of African admixture on the intensity of the response to flu infection (popDR effects). We build individual-wise fold-change (FC) matrices by subtracting non-infected counts from flu-infected counts for each individual (Flu-NI) using weights calculated using the same method as in Harrison et al. 2019. Specifically, given the fold-change entry of: *FC* = *E*^*Flu*^ − *E*^*NI*^, we calculate expected variance of the FC: *σ*^2^(*FC*) = *σ*^2^(*E*^*flu*^) + *σ*^2^(*E*^*NI*^). Within condition weights are: *ω*_*NI*_ = 1/*σ*^2^(*E*^*NI*^) and *ω*_*flu*_ = 1/*σ*^2^(*E*^*flu*^), thus the fold change weight:

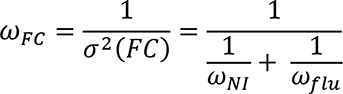

We subset the fold-change matrices to only those features with FDR<.10 for infection effects, since if a feature is not significantly differentially expressed, it cannot be differentially responsive. We then used limma with weights and modeled the effect of admixture on fold changes:

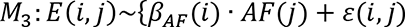

Here, *E*(*i*, *j*) represents the fold change for feature *i* for individual *j* and *β_AF_(i)* signifies the effects of African admixture on feature *i*. For the WGBS, we constructed individual-wise fold-change matrices and subset on CpG sites with FDR < .20 for infection effects. The model used was the same as M3 but including age and batch which for the methylation data are not corrected for *a priori*.

For all data types we performed 1000 permutations obtained by randomly reshuffling admixture estimates in order to estimate FDR, as described previously.

### Calculation of pathway activity scores across individuals

We used the R packages Gene set variation analysis (GSVA), which estimates variation of pathway activity, to calculate individual mean scores for several Hallmark pathways and combinations of gene sets (Hänzelmann, Castelo, and Guinney 2013). The input for GSVA is a matrix of counts and database of gene sets. To apply GSVA to the popDE results, we first obtained all features that are popDE (FDR <.10) in at least one of the conditions. We took the mean of features (peaks or CpG sites) which shared the same closest gene such that there was only 1 value for each gene listed in the Hallmark pathway set. We split the batch and age corrected counts matrix by condition. We then applied the gsva function to each matrix, calculating an individual mean score for each gene set. We used an analogous workflow to calculate individual mean response scores for gene sets using the popDR (FDR<.20) results. The only modification is that we used the fold-change matrices instead of the batch and age corrected counts.

To evaluate the effect of the top SNP and STR on these pathway scores we used an analogous workflow as described above but with the following modifications. We first subset on features associated with genes that were included in any of the immune response pathways tested. We also filtered out the few features that did not have both a SNP and STR associated with the feature. We then obtained the residuals after removing the top SNP and STR effects and followed the same steps as detailed above for the popDR features to calculate ancestry scores. without the effects of the top SNP and STR.

### Elastic net regression to predict transcriptional response based on baseline epigenetic data

We used the glmnet R package (Friedman, Hastie, and Tibshirani 2010) to build an elastic net model to determine if epigenetic features at baseline can predict transcriptional response to flu. Because the number of features is much larger than the number of samples, *glmnet* uses an elastic net penalty to shrink predictor coefficients toward 0. Optimal alpha parameters were identified by grid searching across a range of alphas from 0 (equivalent to ridge regression) to 1 (equivalent to Lasso) by increments of 0.1. We defined the optional alpha as the value that maximized R2 between predicted and true transcriptional response values across samples. We set the regularization parameter lambda to the value that minimized mean-squared error during n-fold internal cross-validation.

To generate predicted transcriptional responses for a given sample, we used a leave-one-out cross-validation approach. Specifically, we separate the samples into training (n-1 individuals) and test (1) samples, where n is the sample size. Training samples were scaled independently of the test sample in each leave-one-out model to avoid bleed-through of information from the test data into the training data. To do so, for each of the datasets, we first quantile normalized the counts data for each feature (or methylation ratios in the case of methylation data) within each sample to a standard normal distribution. Training samples were then separated from the test sample and the normalized counts for each feature (e.g., peak intensity for ATAC-seq data) in the training set were quantile normalized across samples to a standard normal distribution. To predict the transcriptional response in the test sample, we compared read counts for each feature in the test sample to the empirical cumulative distribution function for the training samples (at the same feature) to estimate the quantile in which the training sample methylation ratio fell. The training sample was then assigned the same quantile value from the standard normal distribution using the function *qnorm* in R. A few specific settings were required for the methylation data. First, raw methylation counts were filtered to remove sites that did not have coverage ≥4 sequence reads in at least half of the non-infected or Flu-infected samples and those with 0 variation across all NI samples. Moreover, due to restrictions of the cv.glmnet function, we also removed any CpG sites that had any missing data for any individual, resulting in an input set of 5,528,187 CpG sites.

### Relationship between ancestry-associated gene expression and epigenetic differences

Similar to the analysis described in “Relationship between expression and epigenetic changes in response to infection” we wanted to evaluate if there is a relationship between ancestry-associated gene expression differences and ancestry-associated epigenetic differences. We subset on popDE genes that are higher expressed in individuals with African Ancestry (beta > 0.5, FDR< 0.1) and evaluated how popDE (FDR<0.1) epigenetic differences corresponding to these genes behave. The same is done for popDE features that are higher expressed in primarily European Ancestry individuals (beta < −0.5, FDR < 0.1). A Wilcoxon test was used to determine significance.

### SNP genotype-phenotype association analysis

We used the R package Matrix eQTL (Shabalin 2012) to examine the associations between SNP genotypes and multiple phenotypes of interest (gene expression, chromatin accessibility, DNA methylation levels, and histone marks) in each condition separately. To increase the power to detect cis-QTL, we accounted for unmeasured-surrogate confounders by performing principal component analysis (PCA) on the age and batch corrected expression/peak/methylation matrices. The number of PCs chosen for each data type empirically led to the identification of the largest QTL in each condition and are reported in the table below.

| Analysis | Condition | Regressed PCs |
| --- | --- | --- |
| eQTL | Non-infected | 1 to 4 |
|  | Flu-infected | 1 to 4 |
| caQTL | Non-infected | 1 to 3 |
|  | Flu-infected | 1 to 3 |
| H3K27acQTL | Non-infected | 1 |
|  | Flu-infected | 1 |
| H3K27me3QTL | Non-infected | 1 |
|  | Flu-infected | 1 to 2 |
| H3K4me1QTL | Non-infected | 1 to 2 |
|  | Flu-infected | 1 to 4 |
| H3K4me3QTL | Non-infected | 1 to 2 |
|  | Flu-infected | 1 to 2 |
| meQTL | Non-infected | None |
|  | Flu-infected | None |

Mapping was performed combining individuals in order to increase power, thus, we included the first eigenvector obtained from a PCA on the SNP genotype data as a covariate in our linear model to correct for population structure. For gene expression, chromatin accessibility and histone QTL mapping we used the following model:

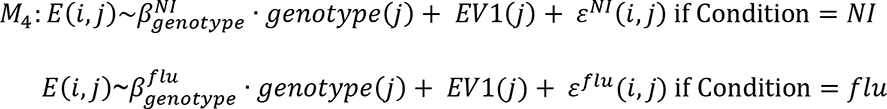

Here, *E*(*i*, *j*) represents the batch and age corrected expression estimates with PCs regressed for feature *i* and individual *j*. EV1 is the first eigenvector derived from the PCA on the SNP genotype data. Local associations (i.e., putative cis QTL) were tested against all SNPs located within the peak or 100Kb upstream and downstream of each peak.

Some modifications were made when performing QTL mapping using methylation proportions due to the nature of the data. First, we quantile normalized across each CpG site using the qqnorm function in R. Since we do not previously account for age and batch as for the other data types, we included mean-centered age and batch as covariates in our model in addition to the first eigenvector obtained from the PCA on the SNP genotype data for the meQTL analysis. Finally, we used a window size of 5kb up and downstream of each CpG site. We did so both to limit the number of tests and because previous studies show that SNPs associated with variation in methylation tend to be located very close to the CpG that they associate with (Banovich et al. 2014; McClay et al. 2015; Huan et al. 2019).

For all data sets, we recorded the strongest association (minimum p-value) for each gene/region/CpG site, which we used as statistical evidence for the presence of at least one QTL for each of the loci tested. We permuted the genotypes ten times, re-performed the linear regressions, and recorded the minimum p-value for each gene/region/CpG site for each permutation. We used the R package qvalue (Storey et al. 2019) to estimate FDR using the permuted p-values as our null expectation. In all cases, we assume that alleles affect phenotype in an additive manner.

In addition to qvalue, we also applied Multivariate Adaptive Shrinkage in R (mashr v0.2.28) (Urbut et al. 2019) to the outputs of the QTL mapping results for each data type independently. For each condition, full Matrix eQTL outputs are loaded (every SNP-feature pair tested). We obtained the effect sizes and the standard error of the effect size, calculated by dividing the beta by the t statistic. For each feature, we chose a single, top *cis-*SNP, defined as the SNP with the lowest pvalue across the two conditions. We recorded the corresponding effect sizes and standard errors of these betas for these top *cis*-SNPs and defined these as our set of “strong” tests. Additionally, we randomly sampled 200,000 rows from all the SNP-feature pairs (including both null and non-null tests). We estimated the null correlation structure using the set of random tests using the zero_Bhat_Shat_reset=2.22044604925031e-16 flag. The data driven covariance matrices were learned using the set of strong tests. We then fit the mash model to the random subset using canonical and data-driven covariance matrices. Lastly, we calculated the posterior summaries for the strong test subset using the fit from the random subset.

### STR calling and filtering

To robustly genotype the highly repetitive STR variants in our samples, we employed the HipSTR algorithm (v0.6.2) (Willems et al. 2017) which accounts for potential sequencing errors of STR introduced through PCR due to the highly repetitive nature of these sequences. Briefly, HipSTR models the PCR stutter noise of the repetitive sequence at each STR locus and determines the most likely STR allele using population-scale data and phased SNP scaffolds. We genotyped a set of 1,504,432 GRCh37 autosomal STRs smaller than 100 using GRCh37.hipstr_reference.bed.gz. We filtered the calls using HipSTR’s supplied script with the recommended thresholds (min-call-qual=0.9, max-call-flank-indel=0.15, max-call-stutter=0.15, -- min-call-allele-bias=−2, min-call-strand-bias=−2) to remove unreliable calls. This resulted in 1,465,954 robustly genotyped STRs.

### STR genotype – phenotype analysis

We use the additive length of both alleles on a STR locus as the genotype to test STR genotype-phenotype association. Each STR locus can have more than three genotypes due to the multiallelic nature of STR length. To ensure that we were only using high quality STR calls, we further filtered the 1,465,954 aforementioned STR set to exclude STRs with call rate < 90% across all samples and STRs with minor allele frequency less than 10%. After filtering, we obtained 442,509 STRs used as input for Matrix-eQTL analysis.

STR-QTL mapping was performed with the same inputs and parameters as the SNP-QTL mapping analysis described above. As with the SNP-QTL analysis, we accounted for unmeasured-surrogate confounders by PCA on the age and batch corrected expression matrices. The number of PCs chosen for each data type empirically led to the identification of the largest QTL in each condition for the STR mapping analysis are reported below.

#### STR PCs reg table

|  |  |  |
| --- | --- | --- |
| eQTL | Non-infected | 1 to 3 |
|  | Flu-infected | 1 to 7 |
| caQTL | Non-infected | 1 to 5 |
|  | Flu-infected | 1 to 5 |
| H3K27acQTL | Non-infected | 1 to 3 |
|  | Flu-infected | 1 to 2 |
| H3K27me3QTL | Non-infected | 1 to 3 |
|  | Flu-infected | 1 to 3 |
| H3K4me1QTL | Non-infected | 1 to 4 |
|  | Flu-infected | 1 to 5 |
| H3K4me3QTL | Non-infected | 1 to 3 |
|  | Flu-infected | 1 to 4 |
| meQTL | Non-infected | None |
|  | Flu-infected | None |

### SNP v. STR analysis

We used a linear model to evaluate the proportion of variance explained (PVE) by the top SNP and top STR on a feature for each genomic phenotype using the R package relaimpo (Grömping 2006). We used a model analogous to QTL mapping adapted to the requirements of relaimpo. The expression values and regressors used to model PVE are closely matched to the ones used for QTL mapping. We used the same batch corrected counts as described in the section “*Infection effects: Infection-Related Differential Effects”*, moving age to the regressor in the model such that the relative importance of the variants could be compared in situations that a feature/gene was only associated with either an SNP-or STR-QTL (i.e., relaimpo requires at least two variables to be included in the model). For DNA methylation data, we used unadjusted, quantile normalized expression value and model it with batch, age, genotype of best associated SNP, and genotype of best associated STR. The relative importance of each regressor to the total variance of the linear model was then reported using the calc.relimp function. We chose to report the lmg relative importance metric as recommended by the package, which outputs the R^2^ of each regressor partitioned by averaging over orders.

### Identification of condition-specific and shared QTL

As in the popDE analysis, we use both qvalue and lfsr to determine if QTL are condition-specific (i.e., only showing an effect in the non-infected or flu-infected conditions) or shared (i.e., showing an effect in both conditions). Specifically, we require SNP/STR-feature pairs to have both FDR <.10 and lfsr < .10 in only one condition to be considered condition specific. If either the SNP-feature or STR-feature pair was found to be condition specific using these thresholds, the feature was classified as condition specific. QTL were classified as shared if FDR < .10 and lfsr <.10 in both conditions for either the SNP or STR.

### Enrichment of TF binding sites among condition-specific SNP QTL

To investigate if condition-specific SNP QTL overlap transcription factor footprints at a significantly higher rate than non-QTL SNP-feature pairs, we used transcription factor footprints detailed in the previous section “Transcription factor activity scores”. Briefly, we overlapped TF footprints with TF motifs and corresponding cluster information. For each data type and condition, we extracted the best SNP for each condition-specific QTL (detailed in “Identification of condition-specific and shared QTL”) and marked if it overlapped with a TF footprint for each of the 200 clusters. This resulted in a matrix containing either 0 (no overlap) or 1 (overlap) for each of the 200 clusters. We collected the same information for the best SNP of all SNP-feature pairs that were not significant (FDR ≥ .10) for that condition, which we used as the background set (for WGBS we randomly sampled 500k from the non-significant pairs). We then performed a logistic regression in R for each cluster to determine if there is a relationship between QTL type (condition-specific v. non-significant SNP-feature pairs) and if the SNP falls within a TF footprint.

### QTL integration across the data types

To determine if SNPs that are QTL (FDR <.10) in one data type are also QTL for other data types we performed the following steps for each data type in each condition: i) collect all significant SNP-gene pairs FDR<.10 (not just the “best” SNPs) for the data type ii) for each of the 6 additional data types, select the top p-value for each feature using the list of significant SNPs for each feature iii) use this top p-value to determine if the SNP is a QTL for the additional data types. By performing this analysis from the perspective of each data type, we ask what percent of QTL are specific to data type or shared among patterns of data types.

To derive a null expectation for the observed overlaps we did the following. Take as an example the expected overlap between eQTLs and the other 6 additional epigenetic QTLs. First, we collected the list of all SNPs that are eQTL (FDR<.10) in the original data. Then, we asked how many of these are also significant for each of the 6 additional data types but using the p-values derived from the permuted results (described in “SNP genotype-phenotype association analysis”). Ultimately, by doing so, we are testing how often we expect to see an overlap between, in this example, an eQTL and other epigenetic QTL just based on the number of association tests performed. These analyses were performed from the perspective of each data type separately.

### Enrichment of TF Binding Sites among shared QTL

To test if shared QTL are more likely to disrupt TF binding than those that are not shared, we modified the QTL integration pipeline to collect *all* p-values for each feature using the list of significant SNPs rather than just the top p-value. We then used each p-value to determine if the SNP is a QTL for the additional data types. For each condition, we took the union across all data types of the SNPs that were shared in 0, 1, etc. data types. For each union set, we marked if each SNP overlapped with any TF footprint. We also collected this information for all SNPs that were tested for QTL mapping to use as the background set. We then performed a logistic regression in R for each union set against the background set to determine if there is a relationship between the number of QTL a SNP is shared across and if the SNP falls within a TF footprint.

### Relationship between epigenetic QTL at baseline and transcriptional response

For each data type separately, we tested the relationship between epigenetic QTL at baseline and transcriptional response. We created a meta genotype for each QTL (using the best SNPs only). We extracted the direction of the effect size from the QTL mapping results at baseline and categorized the 3 possible genotypes as either low/low, low/high, or high/high depending on the direction of the effect. Using the closest gene for the feature, we matched the meta genotype for each individual to the gene expression fold changes quantile normalized to a standard normal distribution. We also tested the relationship between epigenetic QTL and gene expression at baseline and after flu infection using the same steps as above but matching with quantile normalized expression at baseline or after flu infection.

### Enrichment of QTL within popDE features

We tested for an enrichment of QTL among popDE features within each condition. For each condition, we created two vectors: i) a popDE feature vector, where significant features (FDR<.10) were coded as 1 and non-significant features were coded as 0, and ii) a QTL vector, where we extracted the top SNP and STR for each feature and indicated the presence of a significant QTL if either the top SNP or STR feature pair was significant (FDR<.10) by coding a 1. Non-significant variant-feature pairs were coded as 0. A logistic regression was performed using the popDE feature and QTL vectors using glm in R (popDE_status[0,1] ∼ QTL_status[0,1]). The odds ratios were converted to log2 fold enrichments with a 95% confidence interval for plotting.

### Calculating DeltaPVE of Admixture

To evaluate the impact of genetic variation on population differences we calculated ΔPVE of admixture for the significant popDE features (FDR<.10) that have both an associated SNP and STR (i.e., a SNP or STR within the QTL mapping window size for each data type). To calculate ΔPVE of admixture we first calculated the effect of admixture towards the total variance for each batch-corrected feature using the R package relaimpo (Grömping 2006) and the following model:

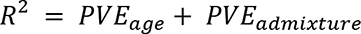

Age was included in the model so that the relative importance of admixture could be compared. We then regressed out the effects of the top SNP and STR:

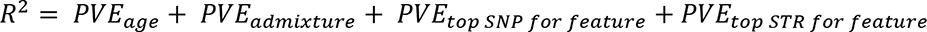

We then calculated the ΔPVE of admixture, which is PVE original - PVE variants regressed / PVE original.

### Calculation of predicted and observed population differences

We estimated the predict cis-genetic population differences across the data types by comparing predicted and observed population differences. For each data type and condition, we extracted significant (FDR <.10) popDE features. We ran a model with inputs analogous to QTL mapping for each data type, but including both SNPs and STRs in the model, in addition to PC1 of the SNP genotype data. Only features that had both a SNP and STR tested were included. For WGBS data, age and batch are also included as regressors since they are not adjusted for in the input file. We regressed out the same number of expression PCs as in “STR PCs reg table”. This resulted in QTL effect sizes with both variants considered. We then computed the predicted “expression” of each feature considering the QTL effect size of the top *cis* SNP for that feature from both the SNP and STR mapping analyses and an individual’s genotype dosage (a vector of 0, 1, or 2 for SNP effect size, and a vector denoting the total length of the STR for the STR analysis). For each feature i, individual j:

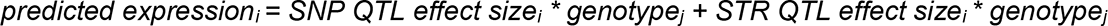

We modeled the predicted expression values using a model analogous to the popDE model (Y ∼ Admixture) in each condition separately since the same features are not always popDE in both conditions. The previously described popDE outputs were used as the observed population differences.

### Evaluating the impact of the top SNP and STR on popDE effects through GSEA

For each data type, we extracted popDE features that were significant in either the NI or flu condition and ran the same model as described in M2 but adding the top SNP and STR for each condition within the model. We used 5 permutations obtained by randomly reshuffling admixture estimates in order to estimate FDR, as described in detail above. We then ran GSEA as described in “GSEA of Infection effects and popDE effects” to compare the size and direction of ancestry-associated effects both before and after regressing out the top SNP and STR for each data type.

### Colocalization analysis

We tested colocalization between the molecular QTLs and 14 well-powered GWAS including 6 unique autoimmune diseases and 4 unique inflammatory diseases (average N: 120132):

| Trait | N | PMID |
| --- | --- | --- |
| Allergy and eczema | 456899 | 29892013 |
| Multiple sclerosis | 115635 | 31604244 |
| Rheumatoid arthritis | 103638 | 24390342 |
| Allergic diseases | 242569 | 29083406 |
| Adult-onset Asthma | 327253 | 30929738 |
| Atopic dermatitis | 40835 | 26482879 |
| IIBDGC – Crohn’s disease | 20883 | 26192919 |
| IIBDGC - Inflammatory bowel disease | 34652 | 26192919 |
| IIBDGC - Ulcerative colitis | 27432 | 26192919 |
| Systemic lupus erythematosus | 10995 | 27399966 |
| Asthma | 142486 | 29273806 |
| Crohn’s disease | 40266 | 28067908 |
| Inflammatory bowel disease | 59957 | 28067908 |
| Ulcerative colitis | 58341 | 28067908 |

We first identified each lead GWAS SNP with P-value below 1e-05 and defined a locus as a 1Mb window centered around the lead GWAS SNP. A GWAS locus was moved from COLOC analysis if its lead SNP overlaps the HLA region (chr6: 25Mb-35Mb). Colocalization test was only performed when the most significant QTL of a feature falls within a defined window (100Kb for RNA-seq, ATAC-seq, H3K27ac, H3K27me3, H3K4me1 and H3K4me3, and 5Kb for WGBS) around the lead GWAS SNP. We used “coloc.signals” function from COLOC package (v5.1.0) with default priors (Giambartolomei et al. 2014; Wallace 2020) We defined colocalization as PP3+PP4 > 0.5 and PP4/(PP3+PP4) > 0.8.

### Imputation of SNPs for heritability analysis

We performed imputation using the same 7,383,243 SNPs for QTL mapping in order to eliminate missing genotypes, as required by the heritability analyses described below. Briefly, we used Genotype harmonizer (Deelen et al. 2014) to harmonize the strand with the 1000 Genomes reference panel. We used SHAPEIT to phase the haplotypes (Loh et al. 2016) prior to imputation with IMPUTE v2 using one phased reference panel (1000 Genomes) (Howie, Donnelly, and Marchini 2009). We imputed each chromosome in 5 MB intervals and used the “pgs_miss” flag to replace only the missing genotypes.

### Fine-mapping molecular QTLs

To better identify likely causal variants, we performed fine-mapping of molecular QTLs using the Bayesian statistical fine-mapping tool SuSiE (Wang et al. 2020). We used SuSiE with individual-level phenotype and genotype data and set the maximal causal variants per region parameter L = 3.

We fine-mapped molecular QTLs with distance-based informative prior inclusion probability, so that a SNP close to a gene or peak would have a higher prior probability of being a causal variant. In specific, we separated molecular QTLs into distance bins, with six distance bins (<500bp, 500bp-1kb, 1kb-2kb, 2kb-5kb, 5kb to 10kb, and 10kb-100kb) for eQTL, caQTL and histone QTLs, and four distance bins (<500bp, 500bp-1kb, 1kb-2kb, and 2-5kb) for methylation QTL. We used the Bayesian statistical tool Torus (Wen 2016) to estimate the enrichment for the distance bins, compute SNP-level priors using the estimated distance enrichment estimates for each locus.

### Heritability and enrichment analysis of GWAS summary statistics using S-LDSC

To partition the heritability of complex traits and estimate heritability enrichment for each type of molecular QTLs we used Stratified LD score regression which assesses how the heritability of a complex trait is partitioned among functional features, while controlling for LD, allele frequency and other baseline features (S-LDSC) (Finucane et al. 2015; Gazal et al. 2017; Bulik-Sullivan et al. 2015) S-LDSC estimates the heritability enrichment as a ratio of the proportion of heritability explained by an annotation divided by the proportion of SNPs in that annotation.

For the enrichment analysis, we constructed a continuous annotation using the posterior inclusion probability (PIP) from SuSiE fine-mapping with distance-based prior. We applied S-LDSC separately for each type of molecular QTL annotations. In our S-LDSC analysis, we adjusted for various baseline annotations of SNPs using a generic baseline LD model, including gene annotations (coding, UTRs, intron, promoter), MAF bins and LD-related annotations. We did not include functional annotations such as enhancer markers in our baseline model, because these annotations are likely correlated with our QTL features of interest and may bias our estimated enrichment.

To estimate heritability explained by molecular QTLs, we constructed a binary annotation containing all SNPs with SNP-level FDR < 10% since the GWAS (same as detailed above in “Colocalization analysis”) used have only been performed on SNPs. We note that the exact values of the heritability estimates may be biased as we have only 35 individuals from a mixture of European and African populations, but the relative heritability estimates should reflect the relative contributions of different molecular QTLs to these complex traits.

### Estimation of the association between genomic marks and immune disease

We used S-PrediXcan (Gamazon et al. 2015) to estimate the association between immune system disorders and the expression of genes and epigenetic marks. S-PrediXcan requires prediction models that describe the association between an aggregate of SNPs and the expression of nearby genomic marks. However, instead of explicitly predicting the genetically determined component of expression, it requires only the summary statistics of GWAS studies to assess the association between a genomic mark and a disorder. We trained a set of prediction models of gene and epigenetic mark expression in both non-infected and flu-infected conditions using the same genotype and phenotype data used for QTL mapping. We used summary statistics from the same 14 GWAS studies previously described to identify the genes and epigenetic marks involved in these disorders. We used the beta and p-value of SNPs from the GWAS summary statistics when available in order to compute the association between the molecular traits and disorders. Otherwise, we used odd ratios. We apply Bonferroni correction to determine the condition and data type specific p-value cut off and identify genes and epigenetic marks that are significantly associated with the immune disorders. We mapped the epigenetic marks to their closest genes using the annotatePeak function of CHIPseeker using the same parameters previously described.

## Supplemental Figures

**Supplementary Figure 1:**
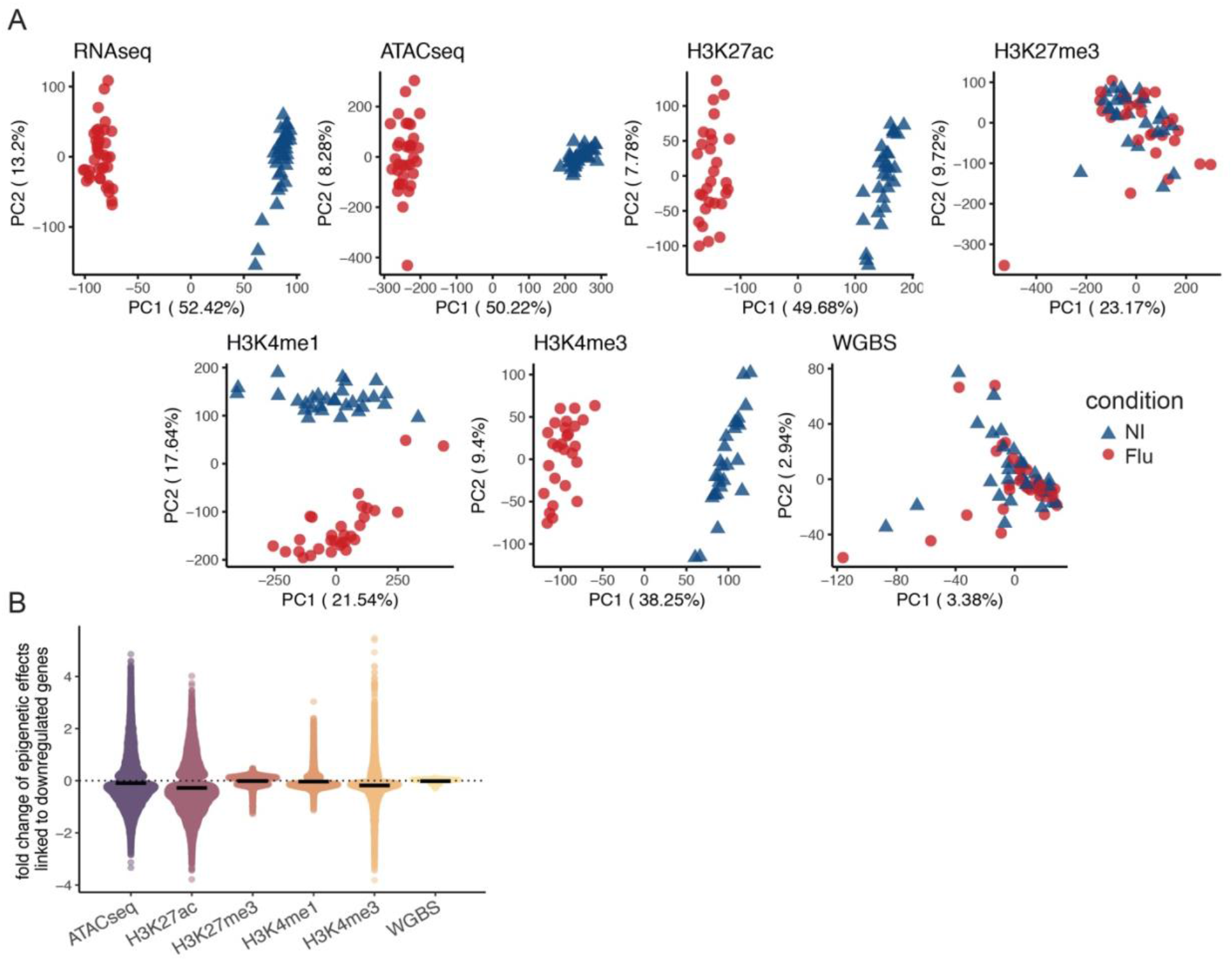
Genome-wide impact of flu infection across regulatory marks. (A) Principal Component Analysis read counts showing for all data types the separation of NI and Flu samples along the two main axes of variation. (B) Distribution depicting the relationship between gene expression changes and epigenetic changes in response to flu infection as seen in Figure 1E but here focusing on epigenetic changes nearby genes that are downregulated in response to infection. Downregulated genes are defined as genes with beta < −0.5 and FDR<.01. Epigenetic changes are those with FDR<.01, except for methylation changes (FDR<.20).

**Supplementary Figure 2:**
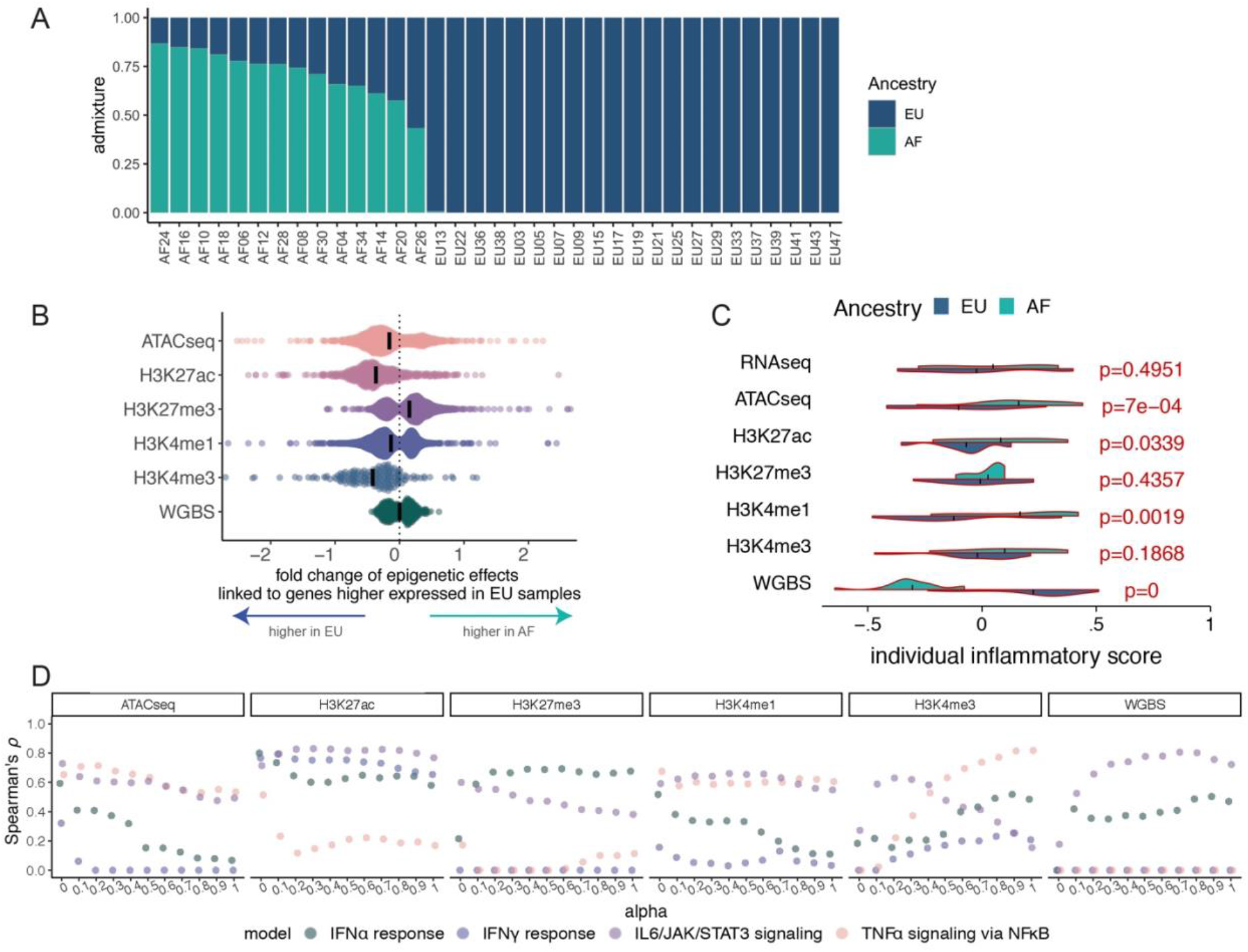
Classification of ancestry-associated differences. (A) Quantitative genetic ancestry proportions partitioned into European (dark blue) and African (turquoise) components for each individual. (B) Distributions of individual mean scores of inflammatory pathways in the flu-infected condition comparable to Figure 2C which shows non-infected condition distributions. A higher score indicates a strong expression of genes or epigenetic marks nearby genes within the Hallmark inflammatory response pathway. (C) Distribution depicting the relationship between popDE genes and popDE epigenetic changes. Genes more highly expressed in individuals with high proportions of European ancestry (fold change < −0.5, FDR< 0.10) are nearby popDE epigenetic regions (FDR <.10) that show increased levels of chromatin accessibility, H3K27ac, H3K4me1 and H3K4me3 in individuals with increased European ancestry levels. Black lines represent means. (D) The distribution of Spearman’s correlation between the predicted and observed mean scores for the various pathways using different alphas.

**Supplementary Figure 3:**
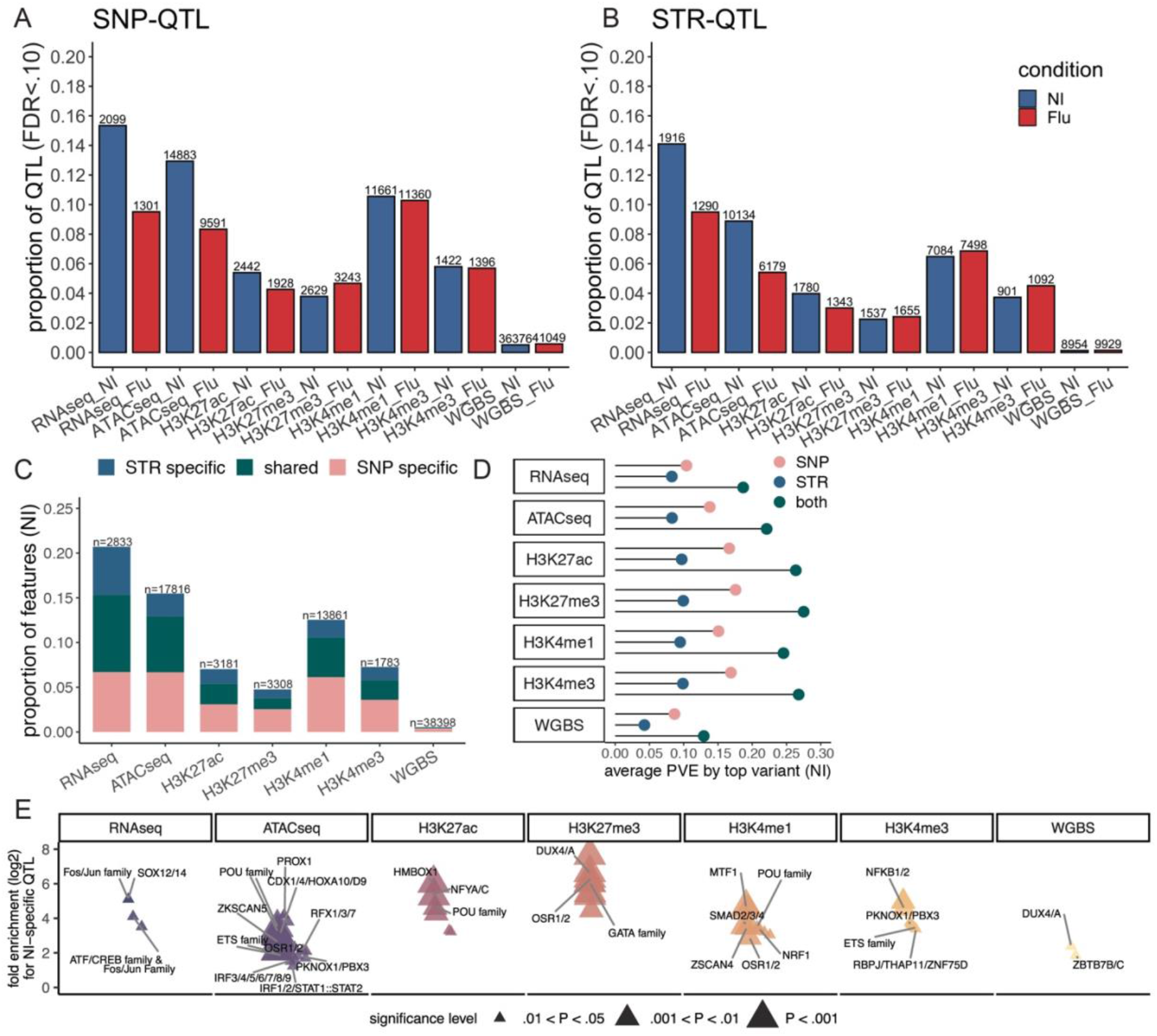
QTL mapping of the different molecular traits. (A) Proportion and number of SNP-QTL at a significance threshold of FDR <.10 in each condition (B) Proportion and number of STR-QTL at a significance threshold of FDR<.10 in each condition. (C) Proportion and number of genes/features associated with at least one SNP or STR QTL in non-infected macrophages. Shared QTL were defined as those genes/features associated with a QTL at an FDR<.10 when performing the QTL mapping against SNPs and STRs separately. SNP- or STR-specific are those only identified as significant (FDR<0.1) against either SNPs or STRs. (D) The mean percent variance explained by the top SNP and STR across all features in the non-infected condition. Both is the sum of the PVE of the top SNP and top STR (E) The enrichment of TF binding sites across non-infected specific SNP-QTL. TF clusters are shown.

**Supplementary Figure 4:**
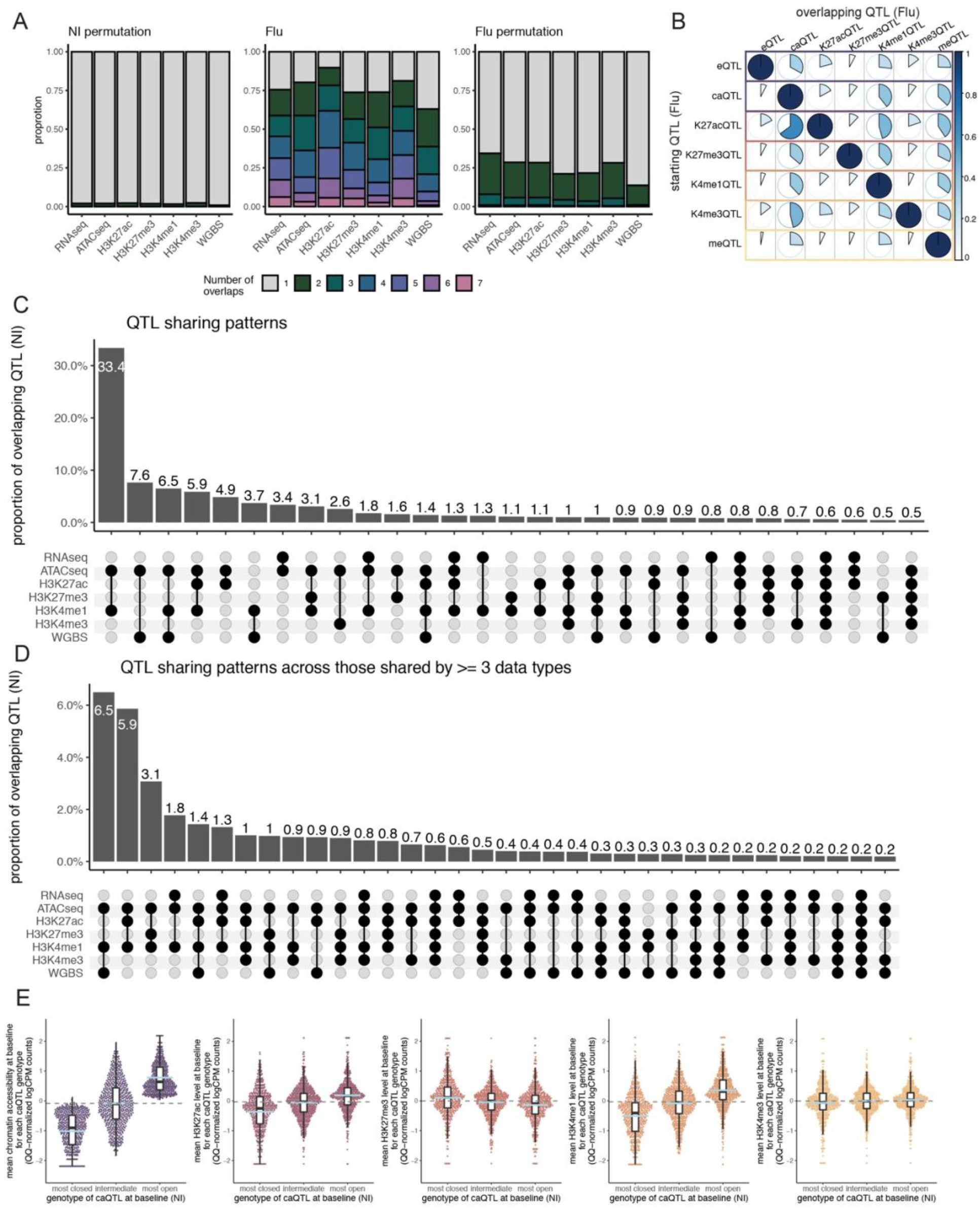
Overlap of QTL across molecular traits. (A) *Left*: The number of overlaps for each QTL type for the permuted analysis in the non-infected condition. More than one overlap indicates the QTL is shared with at least one other datatype. *Center*: The number of overlaps for each QTL type in the flu-infected condition. *Right*: The number of overlaps for each QTL type for the permuted analysis in the flu-infected condition. (B) The percentage of QTL in one data type that are also QTL for another data type in the flu-condition. The starting QTL (rows) are the QTL that are tested for sharing while the overlapping QTL (columns) are the percentage of each starting QTL that are shared with that datatype. The color of each circle corresponds to the percentage of sharing. (C) QTL sharing patterns for those QTL overlapping 2≥ data types) in the non-infected condition. Y axis the proportion of overlapping QTL (i.e., the denominator is the number of QTL that are shared in at least 2 or more data types). (D) QTL sharing patterns for those QTL overlapping 3≥ data types) in the non-infected condition highlighting that caQTL, K4me1 QTL and meQTL are the most commonly shared. The Y axis is the same as described in (C) above. (E) Association between genetically encoded baseline differences in chromatin accessibility and baseline differences in other epigenetic marks. *Left*- Meta caQTL plot (at baseline condition) across caQTLs for accessibility regions associated with up-regulated genes (n=681 caQTLs associated with 506 genes). Individuals with genotypes associated with increased chromatin accessibility also show significantly increased levels of H3K4me1 and H3K27ac (*P*<2.2×10^-16^), and to a lesser extent, a reduction in the repressive mark H3K27me3 (*P*<1.15×10^-10^).

**Supplementary Figure 5:**
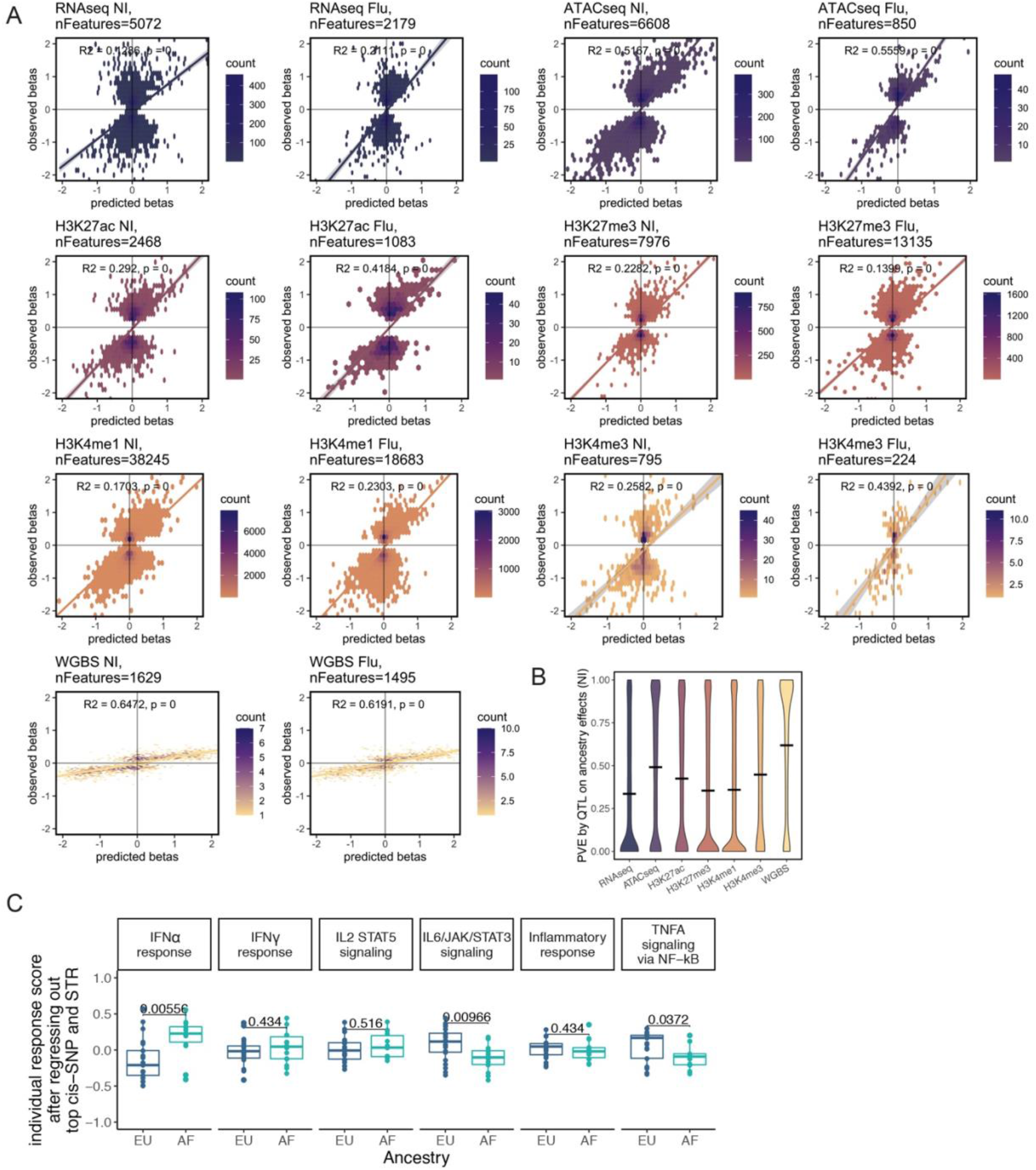
Calculating the contribution of cis-acting regulatory variants to ancestry-associated differences. (A) Correlations between the observed and predicted betas for significant population differentially expressed (popDE) features (FDR<.10) for each of the data types in both conditions (Pearson’s correlation coefficient reported). (B) Boxplot of the ΔPVE of admixture for each feature in each data type in the non-infected condition (flu-infected condition shown in Fig 5C). (C) Boxplots of individual transcriptional response scores after regressing out the effects of the top SNP and STR in each condition for the 6 immune response pathways.

**Supplementary Figure 6:**
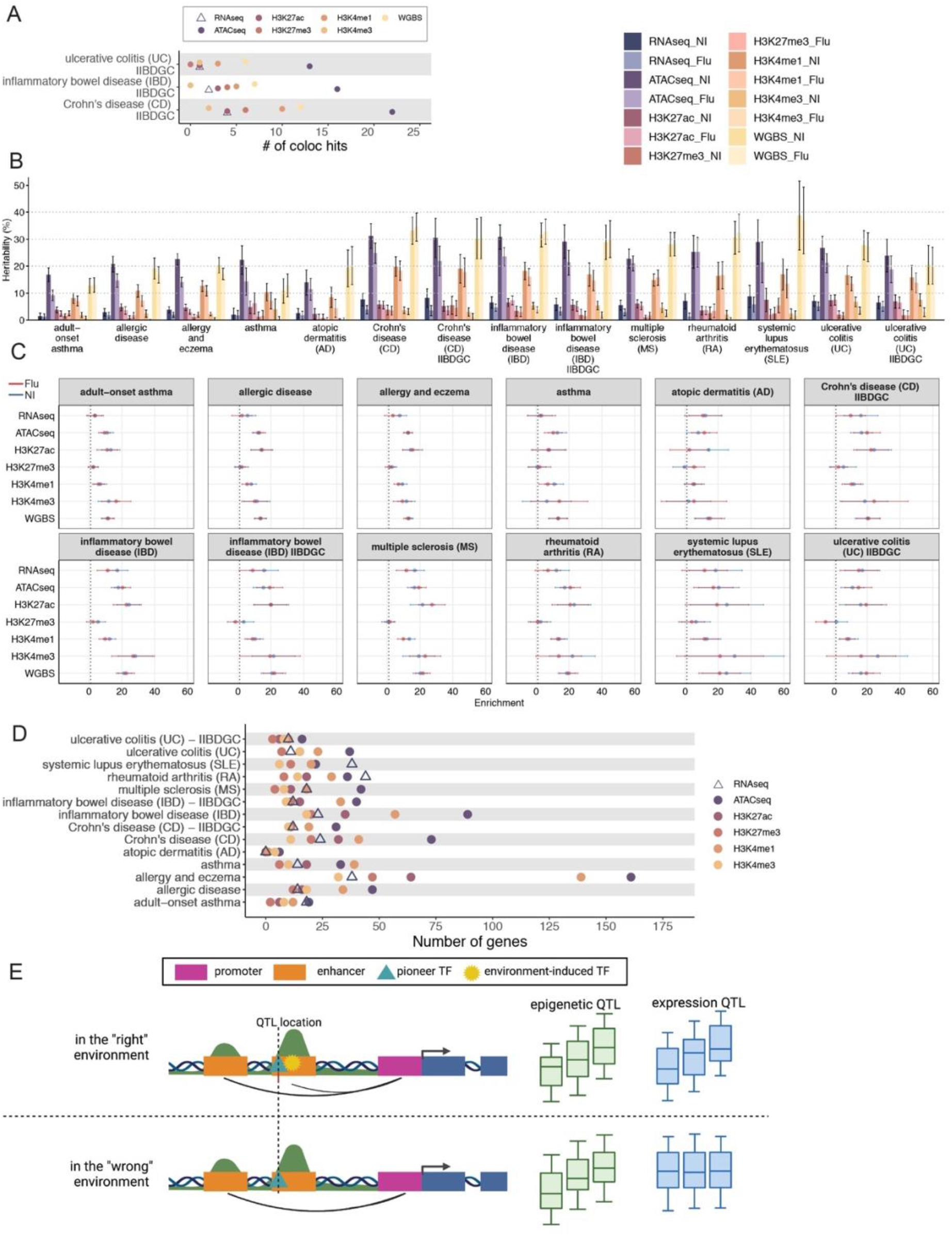
Epigenetic QTLs overlap with genetic variants associated with immune-related diseases. (A) Summary of colocalization results for duplicated immune related diseases (11 diseases were investigated through 14 GWAS). Points represent the number of significant hits defined as PP3+PP4 > 0.5 and PP4/(PP3+PP4) > 0.8 in either condition. (B) Bar plots, with standard error, representing the percent of heritability explained by each of the molecular QTL in all conditions. (C) Heritability enrichment results for all 14 GWAS. A 95% confidence interval is displayed. (D) Summary of PrediXcan results. Each point represents the total number of genes (Bonferroni corrected p=0.05) associated with the disease trait in either condition. A gene is only counted once even if multiple peaks are associated with the gene. (E) Schematic depicting the proposed hypothesis that epigenetic QTL may act as a proxy for genetic variation that under particular environmental conditions has an impact on gene expression levels. Blue boxes represent gene exons and green peaks represent ATACseq peaks. A genetic variant at the QTL location impacts TF binding, such that differential binding of the TF is associated with variation in chromatin accessibility (i.e., an caQTL). If the activity of this enhancer requires the recruitment of an additional TF (here labelled “environment-induced TF”) only induced in response to specific environmental/developmental conditions, the caQTL will not be associated with variation in gene expression levels. Yet, this caQTL will be a proxy for a genetic variant that on the “right environment” will ultimately be associated with an eQTL. Under this model, epigenetic QTLs that colocalize with GWAS variants (but not with eQTLs) can be thought of as a means to identify genetic variants that have an impact on gene expression in a yet unmeasured environment.

## Supplemental Table descriptions

**Table S1.** Description of the samples and libraries generated for this study, related to STAR methods

**Table S2.** List of differentially expressed, accessible and methylated features in response to flu infection, related to Figure 1.

**Table S3:** Transcription Factor activity scores and TF enrichment results in condition specific QTL

**Table S4.** List of population differentially expressed and responsive features, related to Figure 2.

**Table S5.** List of *cis* regulatory QTLs identified in non-infected and flu-infected macrophages using both SNPs and STRs, related to Figure 3.

**Table S6.** QTL integration results, related to Figure 4.

**Table S7.** Colocalization results for 14 immune related GWAS, related to Figure 6.

**Table S8.** Predixcan results for 14 immune related GWAS, related to Figure 6.

Note: Only CpG sites with FDR<.50 in one condition are reported in Tables S2 and S4 and those with FDR <.10 in Table S5 due to file size limitations. Full methylation analysis results available upon request.

